# A rare HSC-derived megakaryocyte progenitor accumulates via enhanced survival and contributes to exacerbated thrombopoiesis upon aging

**DOI:** 10.1101/2024.11.04.621964

**Authors:** Bryce A. Manso, Paloma Medina, Stephanie Smith-Berdan, Alessandra Rodriguez y Baena, Elmira Bachinsky, Lydia Mok, Angela Deguzman, Sarah Beth Avila, Saran Chattopadhyaya, Marcel G.E. Rommel, Vanessa D. Jönsson, E. Camilla Forsberg

**Affiliations:** Institute for the Biology of Stem Cells, University of California, Santa Cruz. Santa Cruz, CA 95064, USA; Baskin Engineering, Biomolecular Engineering, University of California, Santa Cruz. Santa Cruz, CA 95064, USA; Program in Biomedical Science and Engineering, University of California, Santa Cruz. Santa Cruz, CA 95064, USA; Genomics Institute, University of California, Santa Cruz. Santa Cruz, CA 95064, USA; Cellular and Molecular Medicine Program, Johns Hopkins University School of Medicine, Baltimore, MD, USA

## Abstract

Distinct routes of cellular production from hematopoietic stem cells (HSCs) have defined our current view of hematopoiesis. Recently, we challenged classical views of platelet generation, demonstrating that megakaryocyte progenitors (MkPs), and ultimately platelets, can be specified via an alternate and additive route of HSC-direct specification specifically during aging. This “shortcut” pathway generates hyperactive platelets likely to contribute to age-related platelet-mediated morbidities. Here, we used single-cell RNA/CITEseq to demonstrate that these age-unique, non-canonical (nc)MkPs can be prospectively defined and experimentally isolated from wild type mice. Surprisingly, this revealed that a rare population of ncMkPs also exist in young mice. Young and aged ncMkPs are functionally distinct from their canonical (c)MkP counterparts, with aged ncMkPs paradoxically and uniquely exhibiting enhanced survival and platelet generation capacity. We further demonstrate that aged HSCs generate significantly more ncMkPs than their younger counterparts, yet this is accomplished without strict clonal restriction. Together, these findings reveal significant phenotypic, functional, and aging-dependent heterogeneity among the MkP pool and uncover unique features of megakaryopoiesis throughout life, potentially offering cellular and molecular targets for mitigation of age-related adverse thrombotic events.

## Introduction

Aging is associated with remarkable changes to hematopoiesis, resulting in altered blood cell output and function^1–6^. The aging population has an increased risk of developing a host of disorders including cancer^7^, arthritis^8^, diabetes^9^, neurodegeneration^10^, and cardiovascular^11^ diseases; maladies associated with age-related alterations in hematopoiesis^12–15^. One major age-related health concern is platelet-mediated contributions to cardiovascular disease^16^. As such, understanding the cause and consequence of hematopoietic aging is imperative to improve healthy aging. Platelets, anucleate cell fragments, are continuously produced in the bone marrow (BM) by hematopoietic stem and progenitor cells (HSPCs) and function by mediating blood clotting, wound healing, and immune responses^17–19^. With aging, alterations in the number of platelets combined with a lower threshold of activation collectively contribute to platelet-mediated diseases common in the aging population^16,20–30^.

Classically, platelets arise via successive differentiation of hematopoietic stem cells (HSCs) through increasingly restrictive progenitor cell states, including specification into unilineage megakaryocyte progenitors (MkPs, Figure 1A, shaded area) that ultimately mature into megakaryocytes which release platelets into the blood (megakaryopoiesis)^1,2,31^. Recent advances in understanding platelet generation have given rise to new hypotheses of alternative routes of young adult, steady-state platelet specification (reviewed here^1^). Notably, we have definitively demonstrated that an additional, age-related and progressive platelet generation pathway is present during aging^2^ (Figure 1A, dashed red line). Using Flk2-Cre mT/mG (“FlkSwitch”) lineage tracing mice (Supplemental Figure 1A) ^2,32–40^, we discovered that this additional path arises directly from HSCs, bypassing all other progenitors, generating MkPs with unique molecular and functional properties that co-exist with classically-derived MkPs^2^. Importantly, MkPs generated via this additional HSC-direct (tdTomato [Tom]+) pathway give rise to platelets with hyperactive function, likely serving as a key contributor to the onset and progression of platelet-mediated adverse health events that drastically increase upon aging.

**Figure 1.**
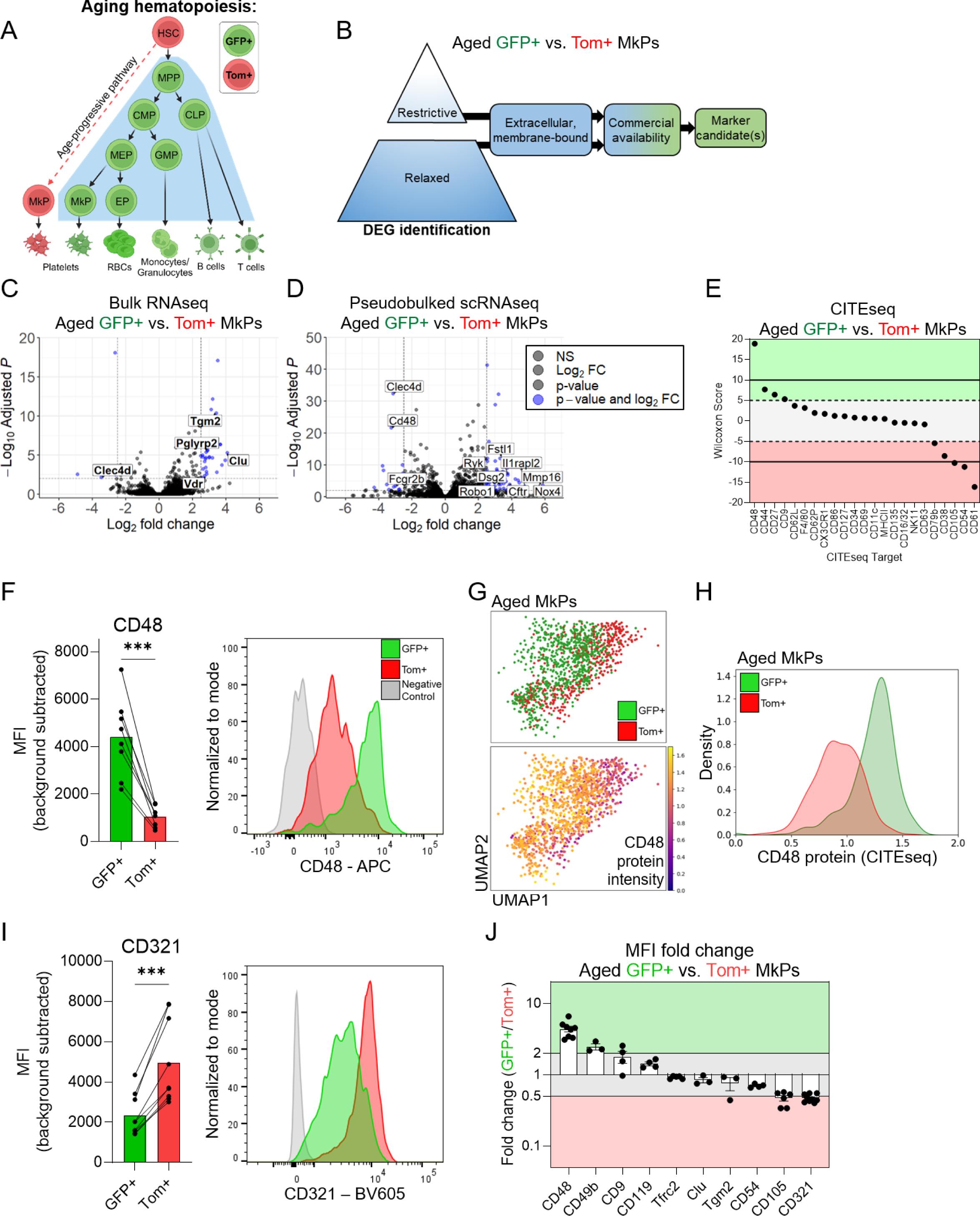
Aged GFP+ and Tom+ MkPs possess unique transcriptomic and proteomic profiles. **A.** Schematic of hematopoiesis in aged FlkSwitch mice^1,2,32^. In this model, HSCs express the tdtomato (Tom) transgene. Upon differentiation to multipotent progenitors (MPPs), recombination mediated by *Flk2*- (*Flt3*)-Cre deletes the Tom transgene and allows irreversible expression of GFP+. Thus, all downstream progenitors and mature cells remain GFP+ for life. Canonical megakaryocytic specification then progresses through a series of increasingly restricted myeloid progenitors including common myeloid progenitors (CMPs), megakaryocyte-erythroid progenitors (MEPs), and megakaryocyte progenitors (MkPs). During aging, a unique and additive population of Tom-expressing MkPs and platelets arise directly from HSCs^2^. Shaded area represents the view of canonical hematopoiesis. Red and green colors represent Tom and GFP expression, respectfully. GMP: granulocyte monocyte progenitor, EP: erythroid progenitor, and CLP: common lymphoid progenitor. **B.** Schematic of candidate marker determination from bulk RNAseq and scRNA/CITEseq. **C-D.** Volcano plots of **C.** bulk and **D.** single cell pseudobulked RNAseq data comparing aged FlkSwitch GFP+ and Tom+ MkPs from Poscablo et al., 2024^2^. Restrictive thresholds are adjusted p-value <0.01 and absolute log_2_ fold change ≥2.5, indicated by blue points. Labeled genes indicate those predicted to encode cell surface proteins that have commercially available flow cytometry compatible antibodies. **C.** 8120 total genes and **D.** 11570 total genes plotted. **E.** CITEseq analysis of aged GFP+ and Tom+ MkPs ran simultaneously with scRNAseq. Dashed lines indicate Wilcoxon Score cutoffs (restrictive cutoffs at 10 and −10, black lines, and relaxed thresholds at 5 and −5, blue lines). **F.** Flow cytometry analysis of CD48 expression (background subtracted median fluorescent intensity [MFI]) among aged FlkSwitch GFP+ and Tom+ MkPs. Each point represents a single mouse with lines connecting cells from the same animal. Paired t-test, ***p<0.001. Example flow cytometry histogram indicating the fluorescence minus one (FMO) control (grey), GFP+ MkPs (green), and Tom+ MkPs (red). n=8 mice across 5 independent experiments. **G.** Uniform Manifold Approximation and Projection (UMAP) of aged GFP+ and Tom+ MkPs from FlkSwitch mice. Top panel indicates MkPs classified as either GFP+ or Tom+ as in Poscablo et al., 2024^2^. Bottom panel demonstrates CD48 CITEseq protein expression intensity across the same cells. **H.** Histogram display of CD48 CITEseq protein expression across aged GFP+ and Tom+ MkPs from the scRNAseq data in **G.** GFP+ MkPs (green) and Tom+ MkPs (red). **I.** Flow cytometry analysis of CD321 expression (background subtracted MFI) among aged FlkSwitch GFP+ and Tom+ MkPs. Each point represents a single mouse with lines connecting cells from the same animal. Paired t-test, ***p<0.001. Example flow cytometry histogram indicating the fluorescence minus one (FMO) control (grey), GFP+ MkPs (green), and Tom+ MkPs (red). n=9 mice across 6 independent experiments. **J.** Summary of cell surface candidates as a measure of fold change in relative protein expression. Horizontal lines indicate two-fold change threshold. Shaded areas indicate if markers are more prevalent on GFP+ (green) or Tom+ (red) aged FlkSwitch MkPs. See also Supplemental Figure 1. Data for CD119 and CD105 from Poscablo et al., 2024^2^.

The molecular features and potential for functional heterogeneity among MkPs remains largely unexplored, particularly during aging. Thus, we sought to determine if the age-unique Tom+ MkPs could be prospectively identified and isolated without the need of genetic reporters and if their enhanced function is rooted in key molecular alterations. We further explore if age-related HSC clonal restriction produces specific MkP subsets, and if so, if we phenotypically identify HSCs posed to generated age-unique ncMkPs.

## Results

### Aged MkPs exhibit distinct molecular and phenotypic heterogeneity

Age-progressive, Tom+ MkPs from old FlkSwitch mice exhibit significant phenotypic, molecular, and functional differences compared to their GFP+ canonical counterparts^2^. Thus, we began by profiling cell surface protein candidates likely to be differentially expressed between aged Tom+ and GFP+ MkPs, with the initial goal of enriching these populations without the need of the FlkSwitch lineage tracing model. To identify candidates, we took a top-down approach using multimodal data of aged GFP+ and Tom+ MkPs from FlkSwitch mice (Figure 1B). We performed in-depth analysis of our published^2,41^ bulk RNA sequencing (RNAseq) and pseudobulked single cell RNAseq (scRNAseq) data for differentially expressed genes (DEGs) using restrictive statistical cutoffs (Figure 1C-D, Table 1). We utilized pseudobulked data to best identify targets that enrich for the MkP subpopulations without biasing for expression outliers. Candidate selection among DEGs was limited to those predicted to code for membrane-bound, extracellular proteins with commonly used and commercially-available flow cytometry antibodies, excluding those used to phenotypically define MkPs (Lin^low^cKit+Sca1-CD150+CD41+, Figure 1B). This approach uncovered five candidates from bulk RNAseq and 11 from scRNAseq (Figure 1C-D, Table 1). Simultaneous with scRNAseq, we generated single cell proteomics via Cellular Indexing of Transcriptomes and Epitopes sequencing (scCITEseq) data of HSPCs from our FlkSwitch mice. Four candidates were identified using a restrictive cutoff (Figure 1E, solid lines, Table 1).

We selected candidates to test with more extreme differences among the restrictive data sets, reasoning that those are most likely to have detectible differences in protein expression by flow cytometry. We initially tested protein expression for the *Tgm2* (transglutaminase 2), *Clu* (clusterin/APO-J), and *Icam1* (CD54) genes by flow cytometry, comparing aged GFP+ and Tom+ MkPs from FlkSwitch mice (Supplemental Figure 1B-D). Of these, only CD54 had significant differences in relative protein levels, yet the magnitude of separation was not compatible with efficient flow cytometry-based cell sorting. *Eng* (endoglin/CD105) and *Ifngr1* (CD119) were also identified as DEGs, which we previously demonstrated have significant differential protein expression^2^.

We also tested *Cd48* (CD48/SLAMF2) as it was identified in the restrictive scRNA/CITEseq data sets. Further, differential CD48 expression was recently reported among young adult MkPs and hypothesized to identify MkPs generated via different routes^42–44^. CD48 demonstrated robust expression among aged GFP+ MkPs with a high degree of separation from aged Tom+ MkPs (Figure 1F). By way of confirmation, overlaying CD48 protein expression (by scCITEseq) on our scRNAseq data further illustrates its differential expression between aged GFP+ and Tom+ MkP subpopulations (Figure 1G-H). Thus, CD48 is a likely candidate for separating the GFP+ and Tom+ MkP subpopulations in old mice.

Flow cytometry cell categorization performs optimally when populations are identified using more than one marker. Thus, we next sought to identify an additional cell surface candidate to pair with CD48 to enrich for GFP+ and Tom+ aged MkPs. Given that strong statistical and log_2_ fold change differences among DEGs do not necessarily reflect detectible differences at the protein level (Supplemental Figure 1B-D), we reasoned that relaxing our inclusion criteria and selecting for DEGs present in both RNAseq datasets would represent likely candidate markers (Supplemental Figure 1E-F, Table 1). We found 23 shared genes between bulk and scRNAseq (Table 2). From these, we selected *Tfr2* (transferrin receptor 2/TFRC2), *Cd9* (CD9/TSPAN29), and *F11r* (CD321/JAM1/JAMA) to test based on the commercial availability of antibodies against their protein products. A relaxation in the scCITEseq criteria similarly confirmed CD9 as a potential candidate (Figure 1E dashed lines, Table 1). We also tested CD49b as it was recently reported to have differential expression among young adult MkPs^42^ and was identified in our relaxed scRNAseq data set (Table 1). Flow cytometry assessment of TFRC2, CD9, and CD49b between aged GFP+ and Tom+ MkPs revealed no clear distinction (Supplemental Figure 1G-I). However, CD321 staining was consistently elevated among aged Tom+ MkPs and distinct from GFP+ MkPs (Figure 1I), representing a second potential candidate.

To determine which candidates are most likely to enable efficient identification and prospective isolation of aged GFP+ and Tom+ MkPs, we evaluated the fold change median fluorescent intensity (MFI) between them for each marker tested (Figure 1J). We reasoned that any robust marker would exhibit consistent staining patterns, have significant separation, and that separation would be a minimum of a two-fold difference. CD48 and CD321 were the only two candidates satisfying all criteria. Importantly, it is advantageous to have one marker that positively selects for GFP+ (CD48) and Tom+ (CD321) aged MkPs.

### CD48 and CD321 coordinately enrich for aged Tom+ MkPs

To assess the combined ability of CD48 and CD321 to enrich for GFP+ and Tom+ MkPs in aged FlkSwitch mice, we first evaluated aged FlkSwitch BM (Figure 2A). Remarkably, GFP+ MkPs were largely CD48+CD321- whereas Tom+ MkPs exhibited the inverse CD48-CD321+ expression pattern. We then performed the reverse approach, first separating aged FlkSwitch MkPs into CD48+CD321- and CD48-CD321+ subpopulations and assessing the GFP+ and Tom+ staining profiles (Figure 2B). Reassuringly, CD48+CD321-MkPs were largely GFP+ and CD48-CD321+ MkPs were primarily Tom+. Thus, CD48 and CD321 selectively enrich for GFP+ and Tom+ MkPs among aged FlkSwitch mice. Importantly, the degree of separation conferred by CD48 and CD321 allow for prospective isolation via flow cytometry, enabling downstream applications. Given that GFP+ MkPs are generated via the canonical differentiation pathway, and Tom+ MkPs via an alternate, age-progressive non-canonical pathway (Figure 1A), we term CD48+CD321- (GFP-like) MkPs as canonical MkPs (cMkPs) and CD48-CD321+ (Tom-like) MkPs as non-canonical MkPs (ncMkPs).

**Figure 2.**
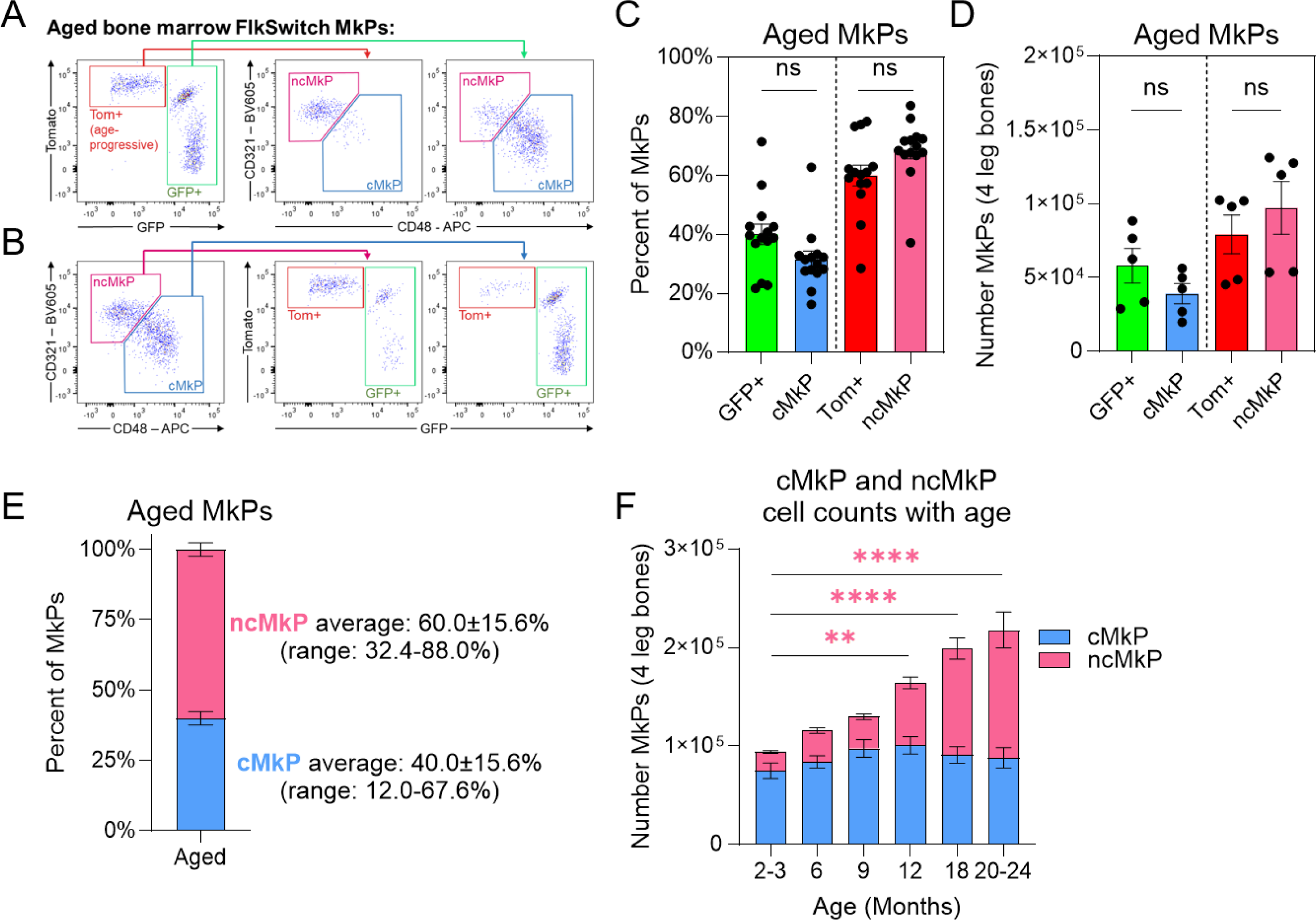
CD48 and CD321 efficiently enrich for GFP+ and Tom+ MkPs in old mice. **A-B.** Example flow cytometry plots demonstrating the efficiency of using CD48 and CD321 in combination to enrich for aged GFP+ and Tom+ MkPs among FlkSwitch mice. **C-D.** The **C.** frequency and **D.** number of aged MkPs defined as GFP+, Tom+, cMkP, or ncMkP. Unpaired t-tests comparing GFP+ to cMkP and Tom+ to ncMkP. **C.** n=14 across 9 independent experiments and **D.** n=5 across 2 independent experiments. **E.** Frequency of cMkP and ncMkP among aged MkPs from wild-type (WT) mice. n=43 across more than 10 independent experiments. **F.** Number of cMkPs and ncMkPs among WT mice at the ages indicated. n=11, 12, 12, 17, 12, and 14 for ages 2-3, 6, 9, 12, 18, and 20-24 months, respectfully across 10 independent experiments. **p<0.01 and ****p<0.0001 evaluated across cMkPs and ncMkPs individually by one-way ANOVA with Dunnett’s multiple comparisons test comparing all populations against the 2-3 month group. Color of the asterisks indicates statistically significant comparisons.

The exceptional enrichment of aged GFP+ and Tom+ MkP populations by CD48 and CD321 is remarkably efficient as the overall frequencies of each related population are similar (comparing GFP+ to cMkPs and Tom+ to ncMkPs, Figure 2C). Quantification of aged FlkSwitch MkPs demonstrate no statistically significant differences between the number of GFP+ MkPs and cMkPs or Tom+ MkPs and ncMkPs (Figure 2D). A minor fraction of cells do not conform to this alternative identification scheme. Indeed, the frequency and number of each subpopulation within individual mice reveals minor, yet statistically significant, differences (Supplemental Figure 2A-B). Overall, CD48 and CD321 possess the stark potential to significantly enrich for aged GFP+ and Tom+ MkPs.

We next sought to evaluate if our cMkP and ncMkP approach recapitulated our previous findings of aged FlkSwitch MkPs. First, we assessed the frequency of aged cMkPs and ncMkPs, which were similar to our previous report^2^ (Figure 2E). The number of ncMkPs increased with advancing age and were statistically significant at 12 months of age (Figure 2F), consistent with the progressive increase in Tom+ MkPs and platelets with age^2^. Importantly, similar to GFP+ MkPs, the number of cMkPs did not change with age. Thus, cMkPs and ncMkPs faithfully recapitulate features of the aged MkP compartment previously only measurable with the FlkSwitch mouse model.

### CD48 protein, but not RNA, expression enriches for aged GFP+ and Tom+ MkPs transcriptomically

To assess the concordance of cMkP and ncMkP enrichment of aged GFP+ and Tom+ MkPs with respect to transcriptional identity, we generated three additional annotations of our scRNA/CITEseq dataset parallel to the original “RNA only, GFP and Tom” annotation^2^ (Figure 3A). First, we subdivided all scRNAseq-annotated MkPs by CD48 protein expression; CD321 was not available for CITEseq (“RNA and CITE,” see Methods and Supplemental Figure 3A). Second, we annotated cMkPs and ncMkPs using only CITEseq (protein) markers, similar to flow cytometry (“CITE only,” Supplemental Figure 3B). As the “CITE only” annotation is the only one that doesn’t rely on RNA-based MkP identification, we assessed if this method accurately captured the transcriptomic MkP cell state. Encouragingly, 93.7% of aged “CITE only” MkPs identified were also classified as MkPs by scRNAseq. Third, using the original “RNA only, GFP and Tom” annotation, we instead divided all scRNAseq-annotated MkPs by RNA expression of *Cd48* and *F11r* (CD321): *Cd48*+*F11r*-cMkPs and *Cd48*-*F11r*+ ncMkPs (“RNA only, *Cd48* and *F11r*,” Supplemental Figure 3C).

**Figure 3.**
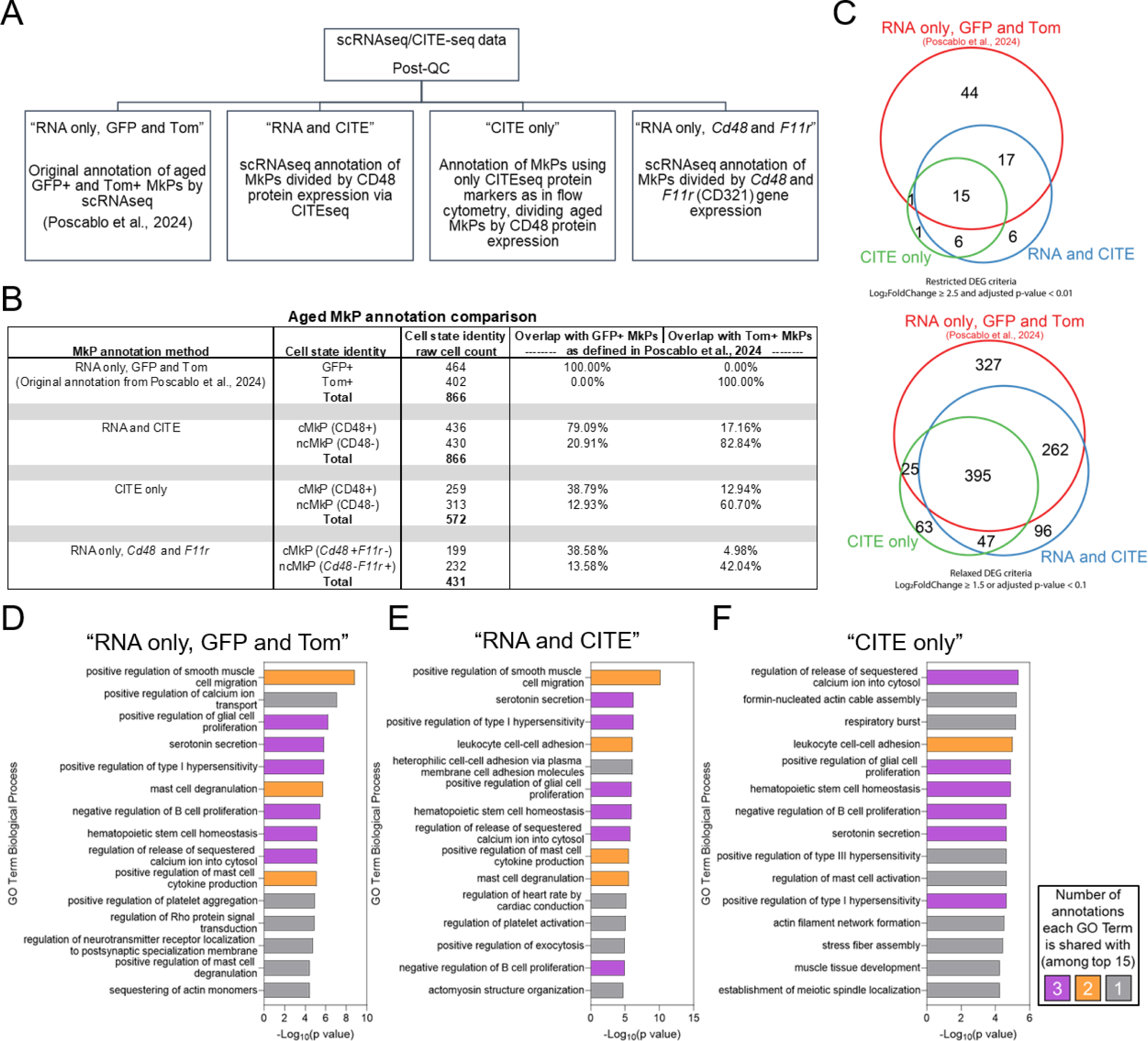
Aged cMkP and ncMkP transcriptomic signatures recapitulate their analogous GFP+ and Tom+ counterparts. **A.** Schematic of additional transcriptomic MkP subtype annotations. **B.** Comparison of the four annotation methods and how they perform compared to the original aged GFP+ and Tom+ MkPs identified via “RNA only, GFP and Tom” annotation^2^. **C.** Venn diagrams comparing the overlap in the numbers of DEGs generated by the three best annotation methods, comparing either aged GFP+ and Tom+ or cMkP and ncMkP populations. Left, DEG comparisons using strict statistical thresholds. Right, DEG comparisons using relaxed statistical thresholds. **D-F.** GO Term Analysis between aged MkP subpopulations across three annotation methods. Top 15 GO Terms displayed. **D.** GFP+ and Tom+ MkPs from the “RNA only, GFP and Tom” annotation. **E.** cMkPs and ncMkPs from the “RNA and CITE” annotation. **F.** cMkPs and ncMkPs from the “CITE only annotation.

Each new annotation identified aged cMkPs and ncMkPs, thus we next sought to compare their transcriptomic fidelity. To do so, we compared each annotation method against the original scRNAseq-based “RNA only, GFP and Tom” annotation that identifies aged Tom+ MkPs from GFP+ MkPs^2^. As each cell in scRNA/CITEseq data is individually barcoded, we compared how often a cell is captured in the “correct” bin across MkP annotations (Figure 3B). That is, of all aged scRNAseq-annotated GFP+ and Tom+ MkPs, how many were also defined as cMkPs and ncMkPs, respectively?

We found that the “RNA and CITE” annotation method performed best compared to the “RNA only, GFP and Tom”. The “CITE only” annotation largely correctly captures GFP+ and Tom+ cells, yet is more conservative in total numbers of cells identified compared to the “RNA only, GFP and Tom” annotation. The “RNA only, *Cd48* and *F11r*” annotation performed poorest, capturing less than 50% of GFP+ or Tom+ MkPs in either of the cMkP or ncMkP bins and was dropped from further assessment. Surprisingly, this may indicate that CD48 and CD321 protein profiles of aged cMkPs and ncMkPs identify them better than their corresponding RNA profiles, an advantage due to the ability to prospectively isolate transcriptomically unique MkPs. Next, we generated restrictive and relaxed DEG lists between aged cMkPs and ncMkPs from the “RNA and CITE” and “CITE only” annotations to assess the amount of overlap with the “RNA only, GFP and Tom” annotation (Figure 3C, Supplemental Table 1). Remarkably, for both restricted and relaxed statistical cutoffs, we find high levels of concordance, indicating that the cMkP and ncMkP cell definitions largely recapitulate aged GFP+ and Tom+ MkPs transcriptomically.

We then subjected the DEG lists from each annotation to GO Term analysis (Figure 3D-F). Given that the small number of DEGs present in restricted DEG lists do not constitute enough statistical power, we utilized the DEGs generated by our relaxed statistical criteria. Importantly, the GO Terms for each annotation returned several overlapping categories, including those related to HSC homeostasis, proliferation, and activation, further supporting the robustness of cMkP and ncMkP designations to recapitulate GFP+ and Tom+ MkPs transcriptomically. Thus, the cMkP and ncMkP paradigm of aged MkPs robustly recapitulates that of their GFP+ and Tom+ counterparts transcriptomically.

### Young mice possess rare ncMkPs

As part of testing CD48 and CD321 to isolate aged GFP+ and Tom+ MkPs, we performed parallel analyses with young mice. MkPs in young BM are classically derived via transition through a Flk2-expressing progenitor cell stage and demonstrate high levels of Cre recombinase-mediated floxing and resulting GFP expression in FlkSwitch mice^2,32^ (Figure 1A). Strikingly, when we assessed cMkPs and ncMkPs in young mice, we detected a rare, yet robust, population of ncMkPs, similar to what others have demonstrated^42–44^ (Figure 2F, 2-3 month age group and Supplemental Figure 4A). Young ncMkPs constitute ∼20% of total MkPs, a significantly smaller frequency than old mice (Supplemental Figure 4B). Unfortunately, the exquisite rarity of young ncMkPs precludes our ability to assess them robustly in our scRNA/CITEseq data. Thus, heterogeneity among MkPs is present in young adult mice, and the pool of ncMkPs specifically expands upon aging (Figure 2F).

### Female mice exhibit modestly reduced effects of aging to the megakaryocytic lineage

The highest efficiency of Cre-recombinase activity is found only in male FlkSwitch mice, thus we have omitted female mice from analyses that rely on detection of GFP and Tom^2^. In contrast, this new cMkP and ncMkP paradigm allows for high fidelity proxy detection and thus we took the opportunity to assess if there are sex dimorphisms among MkP abundance throughout life. Total MkP count in male versus female WT mice were equivalent across six age groups (Supplemental Figure 4C). Male mice had a detectable MkP expansion by 12 months of age whereas female mice lagged behind, only becoming significant around 20 months of age (Supplemental Figure 4D-E). There were no robust differences between male and female mice for cMkPs or ncMkPs (Supplemental Figure 4F, I). Both male and female mice maintained equivalent cMkP numbers throughout life (Supplemental Figure 4G-H) and demonstrated age-related ncMkP expansions with statistically detectable expansion occurring by 12 months of age in male mice and 18 months in females (Supplemental Figure 4J-K). Overall, no major differences in the number of MkPs was found between male and female mice.

Murine platelet counts increase with age via the unique age-progressive pathway^2^. Thus, we also examined total platelet count among male and female wild-type (WT) mice, hypothesizing they would be elevated with aging in both, reflecting the age-related ncMkP expansion. Indeed, platelet counts were elevated upon aging (Supplemental Figure 4L). Interestingly, female mice maintained consistently lower platelet counts compared to their male counterparts (Supplemental Figure 4M). Although age-related expansion of platelet numbers was detected in both sexes, female mice exhibit expansion to a reduced degree (Supplemental Figure 4N-O). Thus, the increase in BM MkPs upon aging results in increased peripheral blood platelets.

### ncMkPs uniquely gain survival, but not proliferative, advantage with age

Given the rare, yet stable, presence of ncMkPs in young mice, we next sought to uncover if MkP subpopulations in both young and old mice were functionally distinct. We first performed in vitro recovery analyses as we previously demonstrated that aged Tom+ MkPs have superior performance^2^. We sorted cMkPs and ncMkPs from young and old WT mice and, following three days of culture, determined the number of viable MkPs remaining. Similar to old Tom+ MkPs, old ncMkPs significantly outperformed all other MkP subsets in recovered viable cell counts (Figure 4A). Young cMkPs, young ncMkPs, and old cMkPs performed equally with no major sex differences (Supplemental Figure 5A-B). Of note, young ncMkPs do not have a functional advantage *in vitro*, highlighting the specific effect of aging on this compartment.

**Figure 4.**
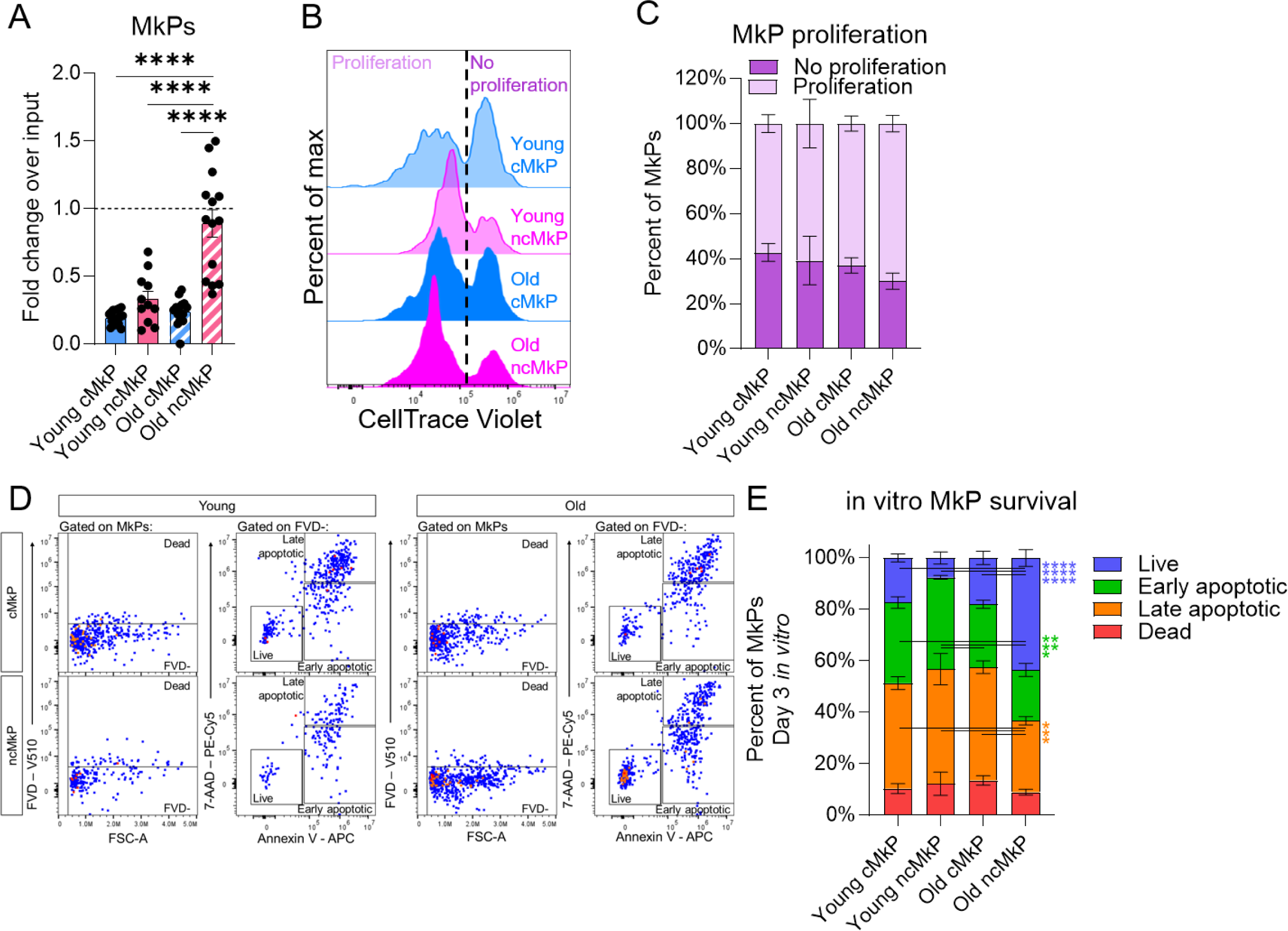
ncMkPs gain a survival advantage with age. **A.** Fold change over input (2000 MkPs) for the number of phenotypic MkPs recovered following three days in culture. Each point represents the average of up to three technical replicates as cell number allowed. n=11-16 mice across 8 independent experiments. ****p<0.0001 by one-way ANOVA and adjusted for multiple comparisons via Tukey’s test. **B.** Histogram representations of CellTrace Violet (CTV) distribution among MkPs cultured in vitro for three days. **C.** Proportion of in vitro cultured MkPs that proliferated or not over the three day culture. n=3-4 across 4 independent experiments. Up to three technical replicates per mouse were averaged together. Statistical non-significance determined via one-way ANOVA adjusted for multiple comparisons via Tukey’s test, no proliferation and proliferation groups tested separately. **D.** Representative flow cytometry plots of in vitro viability determination following three days of culture. Cells were defined as dead if they were fixable viability dye (FVD) positive. Determination of non-dead cell states were made among cells that were FVD negative. **E.** Proportion of viability cell states of three day cultured MkPs. n=3-4 across 4 independent experiments. Up to three technical replicates per mouse were averaged together. *p<0.05, **p<0.01, and ****p<0.0001 by one-way ANOVA adjusted for multiple comparisons via Tukey’s test, each cell state tested separately.

We then sought to uncover the mechanism(s) underlying the enhanced in vitro performance of aged ncMkPs. Proliferation and homeostasis (including cellular survival) were among the top hits revealed by GO Term analyses (Figure 3D-F). Thus, we hypothesized that aged ncMkPs have a proliferative and/or survival advantage. We assessed in vitro proliferation of MkPs using CellTrace Violet (CTV), a fluorescent dye that, upon each cell division, diminishes in intensity via dilution. No significant differences in the frequency of MkPs undergoing at least one cell division were observed across all four subpopulations (Figure 4B-C). We further evaluated the MFI of CTV only among cells that proliferated as a surrogate for the median number of proliferative events (more dye dilution/reduced MFI equates to a greater number of times a given population proliferated). We found statistically significant, yet low magnitude shifts indicating that young cMkPs and old ncMkPs may be equivalently slightly more proliferative than the other populations (Supplemental Figure 5C). Therefore, an increased number of cell divisions could not explain the higher numbers of recovered ncMkPs upon in vitro culture.

Next, we evaluated in vitro MkP survival via combinatorial viability assessment (Figure 4D). This revealed that aged ncMkPs specifically possess a significant survival advantage compared to all other MkPs, which were equivalent to each other (Figure 4E). Viability evaluation was performed simultaneously with CTV, allowing us to determine if MkP viability cell state influences proliferative potential. We stratified the proliferation data across the live, early apoptotic, and late apoptotic cell states (Supplemental Figure 5D-I). Interestingly, nearly all live MkPs proliferated, whereas those destined for apoptosis demonstrate a range of proliferative activity and magnitude. Thus, MkP subset proliferation and survival functions are likely uncoupled (Supplemental Figure 5J).

### ncMkPs are functionally distinct from cMkPs in vivo and demonstrate enhanced function during aging

In vitro regulation of MkPs may differ from in vivo. To assess in vivo regulatory mechanisms of young and old cMkPs and ncMkPs, we first sought to evaluate in vivo proliferation and survival given both the GO Term Analyses and in vitro findings (Figures 3D-F and 4A-E). Analysis of Ki-67, a marker preceding entry into S phase of the cell cycle, from freshly isolated bone marrow (BM) revealed a higher frequency of Ki-67+ cells among young MkP subsets compared to old (Figure 5A). This potentially indicates that young MkPs proliferate and/or enter endomitosis more than aged, or that aged MkPs progress faster through these cycles, thus downregulating Ki-67 more quickly. When sex differences were assessed, slightly more young cMkPs from female mice were found to be Ki-67+ than their male counterparts (Supplemental Figure 6A-B). Additionally, paired analysis between cMkPs and ncMkPs from the same mice revealed no Ki-67 differences between young cMkP and ncMkP populations and a slight, but significant, increase in old ncMkPs compared to cMkPs (Supplemental Figure 6C). Cell cycle classification based on hallmark scRNAseq signatures indicate minor differences between young and old MkP subpopulations, with aged MkPs and, especially aged ncMkPs, possessing fewer cells in a proliferative state (Supplemental Figure 6D). To gain a better understanding of the in vivo proliferative nature of MkPs, we performed 24 hour EdU pulse-chase experiments in young and old WT mice (Figure 5B). Overall, young and old cMkPs and ncMkPs incorporated EdU equivalently, with the exception of slightly diminished incorporation in young ncMkPs, as additionally revealed by paired analysis within individual mice (Supplemental Figure 6E). Thus, in vitro and in vivo analyses of young and aged cMkPs and ncMkPs did not reveal major differences in steady-state proliferation.

**Figure 5.**
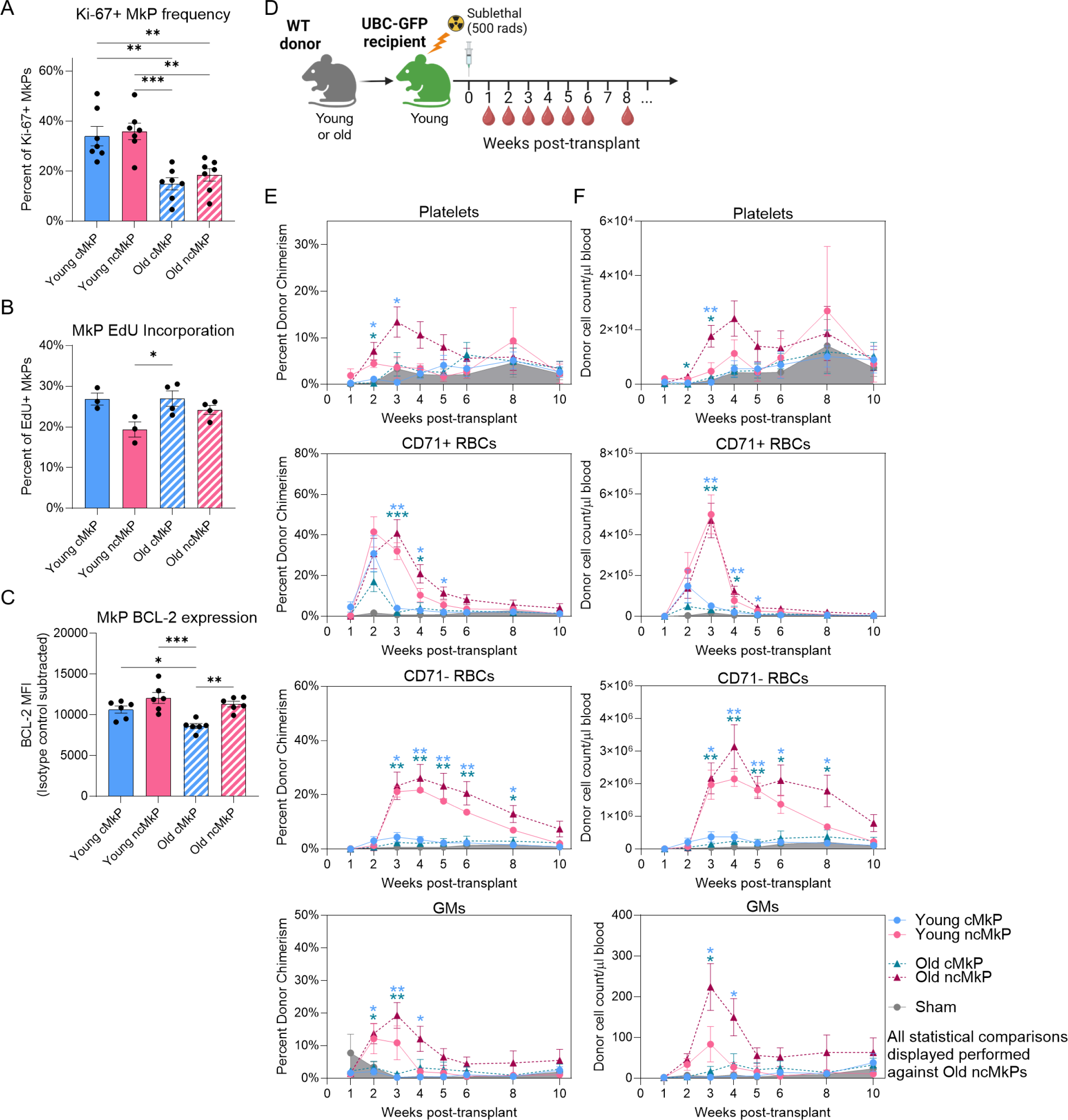
ncMkPs have distinct functional capacity from cMkPs and gain enhanced platelet specification ability upon aging. **A-C.** Flow cytometry analysis of MkP populations from freshly isolated BM. Each point represents an individual mouse, and the frequency or MFI is background-subtracted from an isotype or negative control. *p<0.05, **p<0.01, and ***p<0.001 by one-way ANOVA adjusted for multiple comparisons via Tukey’s test. **A.** Frequency of cells expressing Ki-67. n=7 across 5 independent experiments. **B.** Frequency of cells incorporating EdU 24 hours post-injection. n=3-4 across 4 independent experiments. **C.** Relative BCL-2 abundance as determined by standardized flow cytometry^45^. n=6 across 5 independent experiments. **D.** MkP transplantation experimental design. **E-F.** The **E.** percent donor chimerism and **F.** donor-derived cell number/μl of blood for the indicated populations. Shaded area represents the sham control. Each point represents the mean ± SEM. See supplemental Figure 7A-B, 5 independent experiments. *p<0.05, **p<0.01, and ***p<0.001 by two-way ANOVA adjusted via Dunnett’s multiple comparisons test, comparing each group at each time point to the aged ncMkPs for display simplification.

Aged ncMkPs demonstrated a survival advantage in vitro (Figure 4D-E). Analysis of the bulk and scRNAseq data indicated that gene expression of *Bcl2*, which codes for a major anti-apoptotic protein, was significantly higher in aged Tom+/ncMkPs compared to young and aged GFP+/cMkPs (Supplemental Figure 6F-G). Analysis of freshly isolated MkPs via standardized flow cytometry^45^ revealed high and equivalent levels of BCL-2 protein among all MkP subpopulations, with old cMkPs possessing slightly, but significantly, reduced levels (Figure 5C). Paired analysis demonstrated slightly higher BCL-2 protein levels in ncMkPs compared to cMkPs within each age group (Supplemental Figure 6H). We also assessed MkP viability from freshly isolated BM (Supplemental Figure 6I). Unsurprisingly, the vast majority of cells were highly viable given that dead cells are cleared rapidly in vivo and the likely in situ inhibition of apoptosis by BCL-2. Thus, although difficult to measure in vivo, key molecular features of proliferation and survival among young and aged cMkPs and ncMkPs are grossly similar.

MkPs primarily give rise to megakaryocytes and subsequently platelets in vivo. Given that age-progressive Tom+ MkPs have enhanced platelet producing ability following transplantation^2^, we next tested if aged ncMkP have similarly elevated in vivo performance. We performed transplantations of cMkPs and ncMkPs from young and aged mice into sub-lethally irradiated young recipients and longitudinally assessed blood cell output (Figure 5D-F and Supplemental Figure 7A-D). We expected aged ncMkPs (analogous to aged Tom+ MkPs) to outperform all other populations with respect to platelet generation, subsequently comparing all other conditions to aged ncMkPs. Indeed, aged ncMkPs transiently generated more platelets than young or old cMkPs by measures of donor chimerism and total number of platelets/μl blood. Similar to our in vitro findings, young ncMkPs were indistinguishable from cMkPs (young and old). Thus, aged ncMkPs possess enhanced functional capacity with respect to platelet output following transplantation, recapitulating the output of the analogous age-progressive Tom+ MkPs^2^.

We previously demonstrated that, upon transplantation, MkPs can transiently give rise to erythroid and/or granulocyte/monocyte (GM) lineages^2,41^. Assessment of CD71+ and CD71-erythrocytes revealed that the red blood cell (RBC) output is primarily driven by ncMkPs, regardless of age (Figure 5E-F). Similarly, GM output following MkP transplantation primarily arises from both young and old ncMkPs. As expected, no B or T cell readout was detected (Supplemental Figure 7C-D). Thus, ncMkPs from young and old mice are functionally distinct from their cMkP counterparts in the context of transplantation and, upon aging, gain additional capacity for platelet generation.

### Aged, but not young, HSCs efficiently produce ncMkPs

Young MkPs (GFP+, comprising primarily cMkPs and rare ncMkPs) are specified via the classical path of hematopoiesis as they require transition through a Flk2-expressing cell state typically associated with loss of *bona fide* HSC identity^1,2,46^ (Figure 1A). Alternative hypotheses have been proposed, with CD48^low/−^ MkPs (largely comprising ncMkPs) arising directly from HSCs even in young mice, although our FlkSwitch genetic lineage tracing data is largely incompatible with this view^1^. However, upon aging, a parallel and distinct pathway does arise directly from HSCs to further populate the MkP compartment (Figure 1A)^2^. We therefore sought to uncover if young and old HSCs differentially generate cMkPs and ncMkPs, if age-related changes in the environment influences specification, and if HSCs are clonally restricted to generate either MkP subtype.

We first established an in vitro cell culture assay that is permissive (but not restrictive) to megakaryocytic lineage differentiation. HSCs were sorted from young and old WT mice and cultured for seven days. At the end of the culture period, the total number of live cells produced was equivalent from young and old HSCs (Figure 6A), yet numbers of phenotypic MkPs were elevated over two-fold in aged HSC cultures, analogous to the in vivo state (Figure 6B and 6C, left panels). We then asked if we could identify cMkPs and ncMkPs in vitro using CD48 and CD321. Among phenotypic MkPs, all cells had equivalent CD321 expression (Figure 6C, right panels). In contrast, expression of CD48 recapitulated the pattern observed in vivo: MkPs generated from young HSCs were primarily CD48+ (mean 87.9%, range 75.5-96.7%) whereas MkPs produced by aged HSCs contained distinct populations of CD48+ (mean 70.6%, range 50-91.5%) and CD48- (mean 29.4%, range 8.3-50.0%) MkPs (Figure 6C-D). Indeed, the distribution of in vitro, HSC-derived CD48+ cMkPs and CD48-ncMkPs recapitulated our in vivo observation of an age-dependent increase in ncMkPs (Figure 6D, compare to Supplemental Figure 4B). Further, young and old HSCs generated equivalent numbers of cMkPs, yet aged HSCs produced significantly more ncMkPs (Figure 6E).

**Figure 6.**
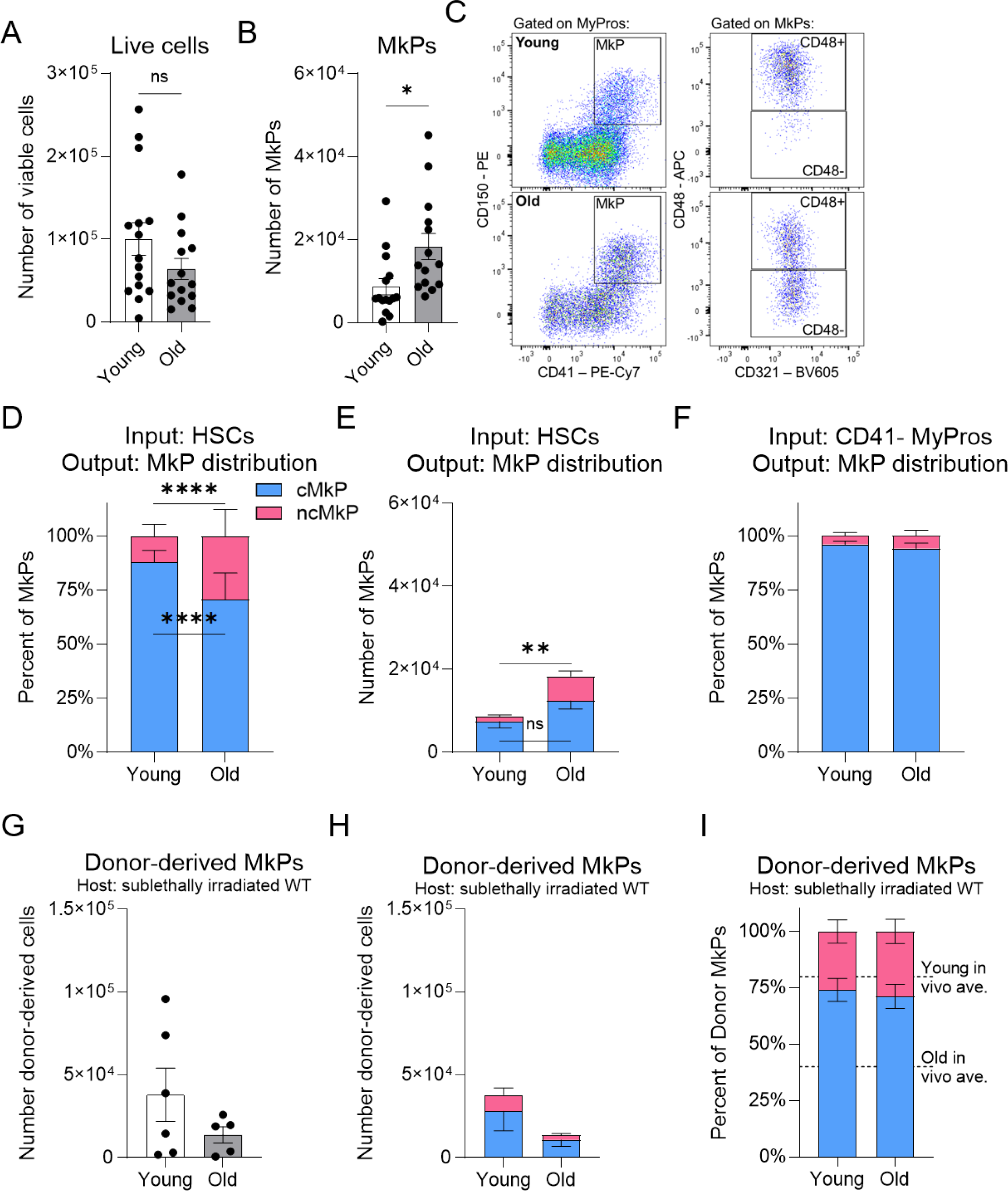
Aged HSCs specifically generate ncMkPs and restore output balance upon exposure to a young environment. **A-B.** Number of **A.** total live cells or **B.** phenotypic MkPs generated following seven days of in vitro HSC culture. Each point represents the average of three technical replicates from individual mice. *p<0.05 by unpaired t-test. n=15 across 9 individual experiments. **C.** Representative flow cytometry plots of in vitro HSC-derived phenotypic MkPs, cMkPs, and ncMkPs. **D-E.** Proportion **D.** and number **E.** of cMkPs and ncMkPs generated by HSCs following seven days of in vitro culture. **p<0.01 and ****p<0.0001 by unpaired t-test. n and number of experiments as in **A-B.** **F.** Frequency of cMkPs and ncMkPs generated by CD41-MyPros following seven days of in vitro culture. Statistical non-significance by unpaired t-test. n=5 across 3 individual experiments. **G-I.** Young or old HSCs were transplanted into young sublethally irradiated (500 Rad) young WT hosts. After 16-20 weeks, BM was analyzed for the **G.** number of donor-derived total MkPs and **H-I.** number and frequency of cMkPs and ncMkPs, respectfully. Each point represents an individual mouse. n=6 young and 5 old from 2 independent experiments. Statistical non-significance by unpaired t-test.

Canonical, but not age-unique, MkP differentiation progresses from HSCs through intermediate progenitor cell states, including myeloid progenitors (MyPros), a heterogeneous collection of myeloid primed cell states (Figure 1A)^1^. To test if myeloid progenitors are capable of producing phenotypic ncMkPs in vitro, we sorted CD41-MyPros (to omit MkPs) and subjected them to the same in vitro assay as HSCs (Figure 6F). Strikingly, both young and old MyPros almost exclusively generated cMkPs, with no difference in distribution.

To test if exposing aged HSCs to a young environment influences MkP subtype specification, we performed HSC transplants from young and old mice into sublethally irradiated young recipients (Figure 6G-I). 16-20 weeks post-transplantation, donor-derived MkPs were equivalently generated from young and old HSCs (Figure 6G). Similarly, young and old donor HSCs produced comparable cMkP and ncMkP numbers (Figure 6H). Importantly, both young and aged HSCs produced similar cMkP:ncMkP ratios, and those ratios reflect young steady-state BM (Figure 6I, compare to Supplemental Figure 4B). Thus, exposure of HSCs to a young environment may overcome the well described decline of aged HSC function with respect to MkP generation.

We next sought to determine if young or aged HSCs are clonally-restricted at the single-cell level to generate cMkPs or ncMkPs (Figure 7A). Platelet lineage bias and restriction has been proposed for subsets of HSCs^1^, and it remains possible that any predisposition is further constrained to specific MkP subpopulations that have different functional capacities. Using our established in vitro system (Figure 6A-F), we sorted individual HSCs from young and old WT mice into single wells of a 96-well plate. Following seven days in culture, the total number of live cells was similar (Figure 7B), mirroring the bulk experimental approach. Among all wells, the number of MkPs generated was variable, with no consistent differences detected between young or old individual HSCs (Figure 7C). Interestingly, this is in contrast to the bulk HSC cultures (compare to Figure 6B). The frequency of single HSCs that generated MkPs (∼40%) was also similar between young and old cells (Figure 7D).

**Figure 7.**
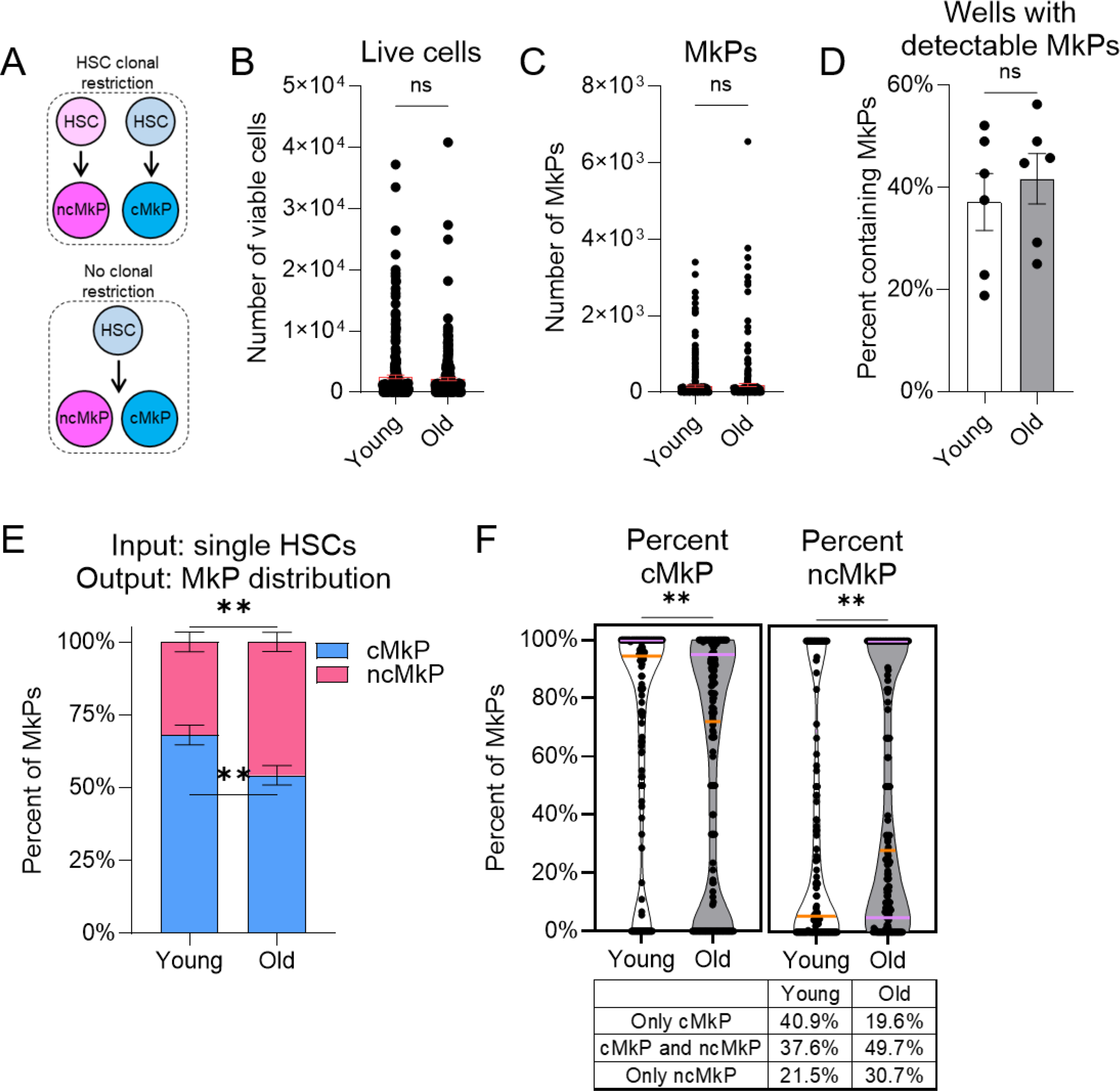
ncMkPs are not exclusively generated by clonally-restricted HSCs. **A.** Model of potential HSC clonal restriction to MkP subtype. **B-C.** Number of **B.** total live cells or **C.** phenotypic MkPs generated following seven days of in vitro single cell HSC culture. Each point represents an individual input HSC. Statistical non-significance by unpaired t-test. For young and old, n=288 individual HSCs each from 5 individual mice across 3 individual experiments. **D.** Among single HSC cultures, frequency of HSCs giving rise to at least one phenotypic MkP. Each point represents an individual mouse, n=6 per age across 4 individual experiments. Statistical non-significance by unpaired t-test. **E-F.** MkP subtype distribution produced by single HSC cultures. **E.** Overall MkP subtype distribution. **F.** Granular analysis of data from **E.** Frequency of cMkP (left) or ncMkP (right) among young and old single-cultured HSCs. Orange line indicates the median whereas the purple lines indicates the quartiles. Each point represents an individually-cultured HSC that gave rise to at least one phenotypic MkP. **p<0.01 by unpaired t-test. Table provides summary of MkP output distribution from individual HSCs. n=149 young and n=163 old across 6 individual experiments.

To assess if MkP subtype generation is clonally restricted to individual HSCs, we calculated the proportion of CD48+ and CD48- in vitro derived MkPs among wells with at least one detectable phenotypic MkP. If HSCs are clonally restricted, we would expect that all MkPs within a well would be exclusively either CD48+ or CD48- (Figure 7A). Conversely, if there was no restriction, we would expect variability in the ratio of CD48+:CD48-MkPs. Overall, single young HSCs generated a higher frequency of cMkPs (and less ncMkPs) than their aged counterparts (Figure 7E), mimicking the bulk data (compare to Figure 6D). A more granular analysis indicates that some young and aged single HSCs gave rise to both subsets whereas others generated only one subtype (Figure 7F). The overall proportionality also indicates the expected age-related shift to ncMkP generation by HSCs. Collectively, a strict clonal restriction model is not compatible with these results.

Given this finding, we sought to predict if specific HSC phenotypes would give rise to cMkPs, ncMkPs, or both. Our single-cell HSC sorting was indexed such that we know the relative expression level of each marker included in the sorting antibody cocktail and which well corresponds to each individual HSC. We then correlated the relative expression of key markers (cKit, CD150, Sca1, CD41, CD48, and CD321) with the number of viable cells, MkPs, and cMkP:ncMkP ratios (Supplemental Figure 8, Supplemental Table 2). Interestingly, no comparison revealed a consistent correlation. Thus, commonly used phenotypic marks among murine HSCs do not predict for in vitro MkP subtype generation, which largely mirrors the in vivo state.

## Discussion

Here, we uncovered new molecular, phenotypic, and functional aspects of age-unique Tom+ MkPs, including faithfully enriching for them without genetic reporters using CD48 and CD321 (Figure 2A-B). In doing so, we also discovered that phenotypic ncMkPs exist as a rare population in young mice that increase over time via the direct HSC “shortcut” path^2^ (Figure 2F). Importantly, we have demonstrated for the first time that both male and female mice have similar megakaryopoiesis aging patterns (Supplemental Figure 4). In contrast to functional decline often associated with aging, old ncMkPs gain enhanced survival (Figure 4E) and, upon transplantation, superior platelet producing capacity (Figure 5E-F). Aged ncMkPs, analogous to the age-progressive Tom+ MkPs, appear to be directly produced by HSCs (Figure 6C-E), but not by MyPros which appear restricted to producing only cMkPs (Figure 6F). We also postulate that the age-related rise of ncMkPs is not wholly clonally restricted at the level of the HSC as some single HSC clones gave rise to both cMkPs and ncMkPs in vitro (Figure 7F). Thus, we have uncovered molecular and functional heterogeneity among young and aged MkPs, supporting and expanding upon previous reports^2,42–44^. We conclude that aged CD48-CD321+ ncMkPs recapitulate the age-progressive, direct HSC-derived Tom+ MkPs.

The cMkP/ncMkP paradigm of aged MkPs is a powerful method by which to investigate distinct pathways of megakaryopoiesis without the need of genetic mouse models. Further, the addition of scCITEseq to our published bulk and scRNAseq of aged FlkSwitch BM HSPCs^2,41^ revealed that stratification of aged MkPs by CD48 protein, but not RNA, best recapitulates the GFP+ and Tom+ subpopulations (Figure 3). Indeed, although a more conservative approach, protein-only identification and separation of aged MkPs largely captures the canonical and age-progressive MkPs. Thus, cell sorting-based experimental approaches utilizing CD48 and CD321 allow for faithful testing of aged GFP+ and Tom+ MkPs.

The idea of molecular and functional heterogeneity among hematopoietic stem and progenitor cell pools is largely accepted^2,6,47,48^. Recently, unique subpopulations of young adult MkPs have been described using only CD48^2,42–44^. We have dramatically expanded upon this by detailing a second marker, CD321, to enhance the separation of these populations. Further, we demonstrate that young and old ncMkPs share some functional similarity in vitro and in vivo, yet aged ncMkPs uniquely gain additional survival and platelet generation capacity.

The observation that young and old ncMkPs possess similar capacity for erythroid and myeloid cell output in a transplantation setting (Figure 5E-F) challenges the hypothesis that phenotypic MkPs as currently defined are unipotent platelet progenitors. However, we do not observe such production in the in situ aging state as old FlkSwitch mice do not possess Tom+ erythroid or myeloid cells^2^. Thus, this additional functional capacity is only induced during transplantation and may reflect the ability of ncMkPs, but not cMkPs, to respond to the acute need for blood cell reconstitution following myeloablation and/or stress. It remains to be seen if perturbations shift ncMkP function to non-platelet production in situ. It is worth noting that cMkPs possess little of this capacity and may therefore be “true” unipotent MkPs. Along these lines, rare young ncMkPs may represent a more quiescent cell state, possibly poised to quickly and transiently respond to stress. Indeed, we find that young ncMkPs are slower to proliferate in vitro (Figure 4C) and in vivo (Figure 5A-B), maintain higher levels of BCL-2 (Figure 5C), and give rise to erythroid and myeloid cells upon acute need in an ablative transplantation scenario.

Young MkPs are GFP+ in FlkSwitch mice^2,32^, and it has been proposed that CD48-MkPs in young mice arise directly from HSCs^42–44^. However, this view is at odds with our FlkSwitch model as the irreversible switch from Tom to GFP happens directly downstream of HSCs (Figure 1A)^1,2,32^. Further, the proportion of GFP+ MkPs in young mice is equivalent to that of Flk2+ MPPs, CMPs, GMPs, and MEPs^1,2,32^, indicating that major intermediate cell states, and not HSCs, are directly upstream of young MkPs. We originally defined ncMkPs in aged mice as being primarily Tom+ (Figures 1-2). Therefore, the surprising finding of a rare population of young ncMkPs may indicate that the age-progressive, HSC-direct (Tom+) MkPs only exist as functionally enhanced MkPs in old mice as their cell surface CD48 and CD321 profile is indistinguishable from that of young ncMkPs. Thus, young mice have two MkP subpopulations; cMkPs and ncMkPs, both of which are GFP+. However, aged mice may have three MkP subpopulations; cMkPs (GFP+), ncMkPs (GFP+), and age-unique (Tom+) MkPs that share CD48 and CD321 expression overlap with ncMkPs. Future investigations will be required to fully deconvolute the phenotypic and functional heterogeneity of MkPs and to identify each subpopulation’s ontogeny.

Because of the prevalence of platelet-related disorders, MkP/platelet ontogeny throughout life is an area under intense investigation^1,2,42–44^. Encouragingly, in vitro-derived MkPs from old HSCs recapitulate the in situ state with respect to increased numbers and altered cMkP/ncMkP frequency compared to young HSCs (Figure 6C-E). It is interesting to note that young and aged CD41-myeloid progenitors made virtually no ncMkPs (Figure 6F), possibly hinting at a more upstream source of this unique cellular pool. Aged ncMkPs (Tom+) arise directly from HSCs^2^; while the cellular origin of young ncMkPs remains elusive, their production is downstream of HSCs and does not appear to involve classical myeloid progenitors. Further, the lack of complete HSC clonal restriction to one MkP subtype indicates that environment or other factors may play a role in the MkP subtype differentiation pathway. Indeed, transplantation of aged HSCs into a young recipient restored young-like ratios of cMkPs and ncMkPs. It is possible that other methods of HSC stratification, such as vWF expression^42^, will better predict MkP subtype generation. Transplantation of clonally restricted vWF+ platelet-biased HSCs results in mutlilineage reconstitution^42^, effectively “resetting” HSC function, similar to our observations (Figure 6G-I). Thus, further investigation is required to uncover the root cause of the age-progressive, non-canonical MkP pathway.

Here, we demonstrate that age-progressive, direct HSC-derived MkPs can be faithfully identified phenotypically and transcriptomically using the surrogate markers CD48 and CD321 to identify the analogous ncMkP (CD48-CD321+). Aged ncMkPs recapitulate their Tom+ counterparts with striking efficiency, enabling widespread experimental application of this paradigm. We also made the surprising discovery that young mice possess very rare ncMkPs that possess enhanced myeloid cell reconstitution capacity upon transplantation compared to their canonical counterparts, yet lack the age-related and paradoxical increased platelet production efficiency of aged ncMkPs. Given that human HSCs gain platelet bias with age^49^, and thrombotic diseases are major health concerns, modulating the number and type of platelets ultimately produced has huge therapeutic potential. Thus, ncMkPs may represent a foundational cellular target to explore for such modulation with the ultimate goal of alleviating patient morbidity and mortality.

## Methods

### Mice

All mice were housed and cared for in an AAALAC-accredited animal facility at the University of California, Santa Cruz (UCSC) and maintained and used under approved IACUC guidelines. Barrier mouse rooms are temperature controlled and on a 12-hour day/night cycle. Mice had ad libitum access to water and food and daily care assessments. Care staff include a trained veterinarian, vivarium manager, and staff. Specific strains used for this study are C57BL/6J (wild-type [WT], Jackson Laboratory #000664), C57BL/6-Tg(UBC-GFP)30Scha/J (UBC-GFP, Jackson Laboratory #004353), and Flk2-Cre x mT/mG mice (FlkSwitch, crossed, bred, and maintained in-house). Aged (20-24 month) WT mice provided by the National Institutes of Health Rodent Ordering System. All mouse ages and numbers indicated in figure legends. WT and UBC-GFP mice were randomized based on sex whereas male FlkSwitch mice were used.

### Mouse tissue collection and processing

When bone marrow was collected, mice were euthanized via CO_2_ inhalation as mandated by the approved UCSC IACUC protocols, followed by cervical dislocation. Hind legs, hips, forelegs, and/or sternum were dissected, cleaned, and immediately placed in staining media (1X PBS [cytiva], 5mM EDTA [Millipore Sigma], and 2% bovine calf serum [BCS, cytiva]) on ice. If cell populations were quantified, a known number of APC Calibrite beads (BD Biosciences) were added immediately prior to processing. Bones were processed either by crushing via mortar and pestle or spin-isolated in staining media^50^. Following cell release from bones, cells were passed through a 70μm filter, and centrifuged for 5 minutes at 300xg. Peripheral blood was collected via nicking the tail vein. A known volume of blood was immediately added to staining media containing a known number of APC Calibrite beads (BD Biosciences) and appropriate antibodies for flow cytometry analysis (see Flow Cytometry methods). Mouse ages as indicated in figures, and young mice are 8-12 weeks old whereas old/aged mice are 20-24 months.

### Hematopoietic stem and progenitor cell enrichment

Hematopoietic stem and progenitor cells were enriched from freshly isolated whole BM via positive magnetic selection. Cells were suspended in 0.5-1.0mL staining media and incubated with 2μl/bone anti-CD117 (cKit) magnetic beads (Miltenyi Biotec) for 30 minutes on ice, mixing at the 15 minute mark. Positive selection via LS columns (Miltenyi Biotec) was performed according to manufacturer’s instructions. The cKit-enriched cell fraction was then washed in staining media and used immediately.

### Flow Cytometry and cell sorting

#### Whole or cKit-enriched BM

Cells to be stained were aliquoted into either 5mL polystyrene tubes or individual wells of a 96-well U-bottom plate. Cells were then blocked for a minimum of 10 minutes on ice in a solution of staining media with 5% each normal mouse and rat serums (Invitrogen). If the experimental design used unconjugated mouse-derived antibodies (and anti-mouse secondary), only those were used first, followed by blocking. The appropriate antibody cocktail was then added directly to cells plus block, and incubated for 30 minutes on ice in the dark. When CXCR4 staining was performed, cells were instead incubated at room temperature for 45 minutes in the dark. Cells were then washed and either stained with streptavidin secondary for 20 minutes on ice in the dark or resuspended for use. Cells were thoroughly washed with staining media between each staining step. Immediately prior to acquisition, 3 drops of 1:5000 diluted propidium iodide (PI, MP Biologicals) was added and cells were mixed. Where indicated, negative controls (fluorescent minus one or isotype) were employed.

#### Peripheral blood

The primary antibody cocktail was prepared in staining media together with a known number of APC Calibrite beads (BD Biosciences) and placed on ice. A known volume of blood was collected (typically 10μl) and immediately mixed well in the antibody/bead solution. Cells were then treated as described for BM above. Following completion of staining, ∼10% of the homogenized and stained blood was aliquoted and diluted for whole blood (platelet and erythrocyte) analysis. The remaining sample was subjected to erythrocyte lysis via ammonium-chloride-potassium (ACK, 0.15M NH_4_Cl, 1mM KHCO_3_, and 0.1mM Na_2_EDTA in diH_2_O, pH 7.2) incubation. Cells were washed, pelleted, and resuspended in 1X ACK at room temperature for 5 minutes. Cells were then washed with staining media. Immediately prior to acquisition, 3 drops of PI solution were added followed by cell mixing.

#### In vitro and post-migration analysis

At termination of in vitro and migration cultures, a known number of APC Calibrite beads (BD Biosciences) was added to each well. Plates were then centrifuged for 5 minutes at 300xg (∼1250 RPM). Supernatant was manually removed via pipetting, and samples were blocked and stained as described above. To preserve rare cell numbers, cells were not washed but resuspended in 1:5000 diluted PI following completion of staining.

#### CellTrace Violet (CTV) staining

Freshly isolated BM from young and aged mice were prepared and cKit-enriched as described above. Following antibody staining, the enriched fraction was then labeled with CTV following manufacturer’s instructions. Labeled cells were then isolated via fluorescence active cell sorting (FACS), selecting for strong and uniform CTV signal by double-sorting each population.

#### Flow cytometric viability assessment

Freshly isolated BM or in vitro culture endpoint samples were stained with antibodies for analysis as described above, excepting use of the Annexin V binding buffer (BioLegend) instead of staining media. Ghost Dye™ Violet 510 fixable viability dye (FVD, Cytek) was included simultaneously with antibody staining at a final concentration of 1:1000. Following staining, Annexin V and 7-AAD (BioLegend) was added for 15 minutes at room temperature in the dark as per manufacturer’s instructions. Annexin V binding buffer was then added and samples were immediately acquired.

#### Ki-67 staining

Cell-surface antibodies were used as described above. Cells were then fixed, permeabilized, and stained with the FoxP3 kit (BioLegend) per manufacturer’s instructions. An isotype control of equal protein amount was used to determine percent Ki-67 positive cells. Following completion of staining, cells were resuspended in 1% PFA (Electron Microscopy Sciences) in 1X PBS for analysis.

#### In vivo 5-ethynyl 2’-deoxyuridine (EdU) incorporation and detection

EdU was dissolved in 1X PBS and injected intra-peritoneal (i.p.) at a concentration of 50mg/kg mouse weight. 24 hours post-injection, mice were sacrificed and BM prepared as described above. Detection of EdU was performed via the Click-iT™ Plus EdU Flow Cytometry Assay Kit (Invitrogen) per manufacturer’s instructions. Negative control mice were EdU-injected, but did not undergo the Alexa Fluor™ 647 labeling step.

#### BCL-2 staining

Cell-surface antibodies were used as described above. Cells were then fixed, permeabilized, and stained with the True-Nuclear kit (BioLegend) per manufacturer’s instructions. An isotype control of equal protein amount was used to determine relative BCL-2 expression following standardized flow cytometry approaches^45^. Following completion of staining, cells were resuspended in 1% PFA (Electron Microscopy Sciences) in 1X PBS for analysis.

#### Cytometers

FACS was performed via a BD Biosciences FACSAria IIu. Cell analysis was similarly performed on the FACSAria or Cytoflex LX (Beckman Coulter). All flow cytometry and FACS data was analyzed via FlowJo (BD Biosciences). The FlowJo plugin IndexSort was used for analysis of single sorted HSCs.

#### Phenotypic cellular definitions and example flow cytometry staining

Supplemental Figure 9 details all cellular definitions used illustrates representative flow cytometry data for each cell type and experimental approach.

### Single cell RNAseq analyses

We performed pseudobulk differential expression analysis using the Python3 implementation of DESeq2^51,52^ as previously performed^2^. Briefly, we restructured the single-cell dataset into sample-level datasets. We extracted the sample and cell type clusters for young GFP+, old GFP+, and old Tom+ MkPs and aggregated our scRNAseq data into pseudobulk samples by summing the counts of individual cells within each biosample and condition. We ran DESeq2 on the pseudobulked count matrix to generate a list of differentially expressed genes between two conditions of interest. We also computed the baseMean of each gene within a condition by taking the mean normalized counts of a gene across cells within that condition. The DEG lists were then further filtered by removing mitochondrial and ribosomal gene transcripts, as well as Malat1 transcripts, to reduce technical bias, as well as those whose baseMean values were less than 5 in both conditions or had no calculated adjusted p-value. We subsequently conducted a GO enrichment analysis using DEGs generated from pseudobulk DEseq2 output using GOATOOLS. GO Terms reported required at least 20% DEG representation for each GO Term. The scores of cell cycle phases were calculated using the Scanpy function ‘score_genes_cell_cycle’ on the basis of canonical markers^53^.

### Cellular indexing of transcriptomes and epitopes sequencing (CITEseq)

#### Sample preparation

CITEseq via the 10X Genomics platform was performed simultaneously with our previously published scRNAseq data of young and aged FlkSwitch mice^2^. Briefly, cKit-enriched BM was sorted into CD150+ and CD150-fractions and incubated with CITEseq antibodies (BioLegend) per manufacturer’s instructions. CD321 was not included as it was not available. All other steps as described in Poscablo et al., 2024^2^. Chromium Next GEM Single Cell 3’ v3.1 (Dual Index) reagents and protocols utilized as specified. Sequencing depth of 5,000 read pairs per cell was performed.

#### CITEseq data preparation and QC

Cells expressing fewer than 200 genes were excluded from the analysis. Cells were further filtered based on surface protein marker expression; specifically, double-negative cells for CD117 antibody-derived tag (ADT) and *Kit* RNA were excluded, removing 13 cells from the dataset. To normalize the CITEseq data, we implemented a centered log-ratio (CLR) transformation for each cell^54^. For each cell’s ADT count vector, we first computed the natural logarithm of the counts, offset by one, for all non-zero elements. The sum of these log-transformed values was then divided by the total number of features, and the exponential of this mean was used to normalize the original counts. Finally, a log transformation was applied to the resulting normalized values to stabilize variance across features.

#### Differential protein expression analysis

We quantitatively compared ADTs generated by CITEseq using the sc.rank_genes_group function that uses the t-test_overestim_var method in the Scanpy python package. Although included in the analysis, lineage (CD3, CD4, CD8a, B220, Gr1, Ter119, and CD11b) and MkP-defining (CD117/cKit, Sca1, CD150, CD41) markers were excluded from the display of Figure 1E.

#### Additional annotations

We generated three annotations of interest:

1. RNA annotated MkPs^2^ further split into cMkP (CD48+) and ncMkP (Cd48-) via CITEseq CD48 expression (Supplemental Figure 3A).
2. CITEseq-only annotation of cMkPs (CD48+) and ncMkPs (CD48-), mimicking flow cytometric-based cell subsetting (Supplemental Figure 3B).
3. RNA-based classification using *Cd48* and *F11r* (CD321) expression utilizing defined MkPs^2^. Cells were evaluated for *Cd48* and *F11r* expression and classified as either *Cd48*+*F11r*- or *Cd48*-*F11r*+ (Supplemental Figure 3C). We excluded 98 cells that had both *Cd48* and *F11r* expression, as well as 481 cells that had no *Cd48* or *F11r* expression.

Venn Diagrams of annotation version comparisons generated via the R package Vennerable, with input as described in the text and figure legends.

Each annotation set was further divided by sample age (young and old). RNA only annotations were previously done by Poscablo et al., 2024 where hematopoietic cell states were annotated using scRNAseq data. For CD48 protein call of positivity (annotations (1) and (2) above), we applied a two-component Gaussian Mixture Model (GMM) via the GaussianMixture function in Scikit-learn^55^ to CD48 protein expression among cells annotated as MkP via RNA expression. From the inferred mean values of the two components, we derived the CD48 positive and negative group threshold as the mean of these components: 1.1. For annotation (2), we manually derived thresholds for CITEseq proteins of interest to annotate MkPs via protein expression. MyPros were annotated at expression values of Sca-1 < 0.71 and CD117 > 1. We subsequently annotated these myeloid progenitor cells as MkPs if they had CD150 > 0.45 and CD41 > 0.6 expression. Lastly, these MkP CITE annotated cells were classified as CD48+/− by applying the previously determined threshold of 1.1.

### Selection of cell surface MkP candidates via bulk RNAseq and scRNA/CITEseq

Bulk RNAseq comparisons of aged GFP+ and Tom+ MkPs as described previously^2^ were deeply analyzed. First, DEG lists were filtered to move any genes with an average read count of 50 or less across both cell states, as well as those with no computed adjusted p-value. Next, we applied two possible thresholds to identify genes with the largest differences; a strict threshold consisting of genes with a Log2FoldChange ≥ 2.5 and adjusted p-value <0.01 and a relaxed threshold of Log2FoldChange ≥ 1.5 or adjusted p-value <0.1. Finally, genes were further filtered to select for those predicted to be membrane-bound, extracellularly expressed, and with commercially available antibodies against an individual gene’s protein product. The scRNAseq data was similarly treated (see above), with the only exception of removing any genes with an average read count of 5 or less across both cell states. Volcano plots of DEGs were generated via the EnhancedVolcano package in R, labels assigned as detailed in figure legends.

CITEseq candidates were assessed via Wilcoxon rank score comparing aged GFP+ and Tom+ MkPs. Protein targets of mature lineage cells (CD3, CD4, CD5, CD8a, CD11b, Gr1, B220, and Ter119) were not included in the analysis. Similarly, protein targets used to define the MkP cell state (cKit/CD117, Sca-1, CD150, and CD41) were not included. A restrictive Wilcoxon score cutoff of 10 and −10 was applied, as well as a relaxed threshold of 5 and −5.

### Bulk in vitro cultures

HSCs, CD41-MyPros, cMkPs, and ncMkPs were isolated via FACS from fresh BM as described above. All populations were double-sorted to ensure purity. Sorted cells were cultured in a 96-well U-bottom plate and incubated at 37°C, 5% CO_2_. 200 HSCs, 2000 CD41-MyPros, 2000 cMkPs, or 2000 ncMkPs were plated per well, and up to three replicates were setup per mouse and cell type. HSCs and CD41-MyPros were cultured for seven days in IMDM plus GlutaMAX™ (Gibco) containing 20% fetal bovine serum (FBS, Atlas Biologicals), 1X Primocin (InvivoGen), 1X non-essential amino acids (Gibco), and recombinant mouse thrombopoietin (TPO, 50 ng/ml), stem cell factor (SCF, 20 ng/ml), IL-6 (20 ng/ml), IL-3 (10 ng/ml), and IL-11 (20 ng/ml, all Peprotech). MkP subpopulations were cultured for three days in IMDM plus GlutaMAX™ containing 10% FBS, 1X Primocin, 1X non-essential amino acids, and recombinant mouse TPO (20 ng/ml), SCF (50 ng/ml), IL-6 (20 ng/ml), and IL-3 (5 ng/ml) as previously described^2,41^.

### Indexed single cell HSC in vitro cultures

Single HSCs were prepared, isolated, cultured, and analyzed as described above, with the exception of using the BD FACSDiva single-cell sorting software module per manufacturer’s directions, with the index sorting option selected.

### MkP transplantations

cMkPs and ncMkPs from young and aged mice were freshly isolated via FACS from cKit-enriched BM as described above. Populations were sorted twice to ensure specificity. For each individual transplant, 22,000 MkPs were injected retro-orbital in 50μl 1X HBSS (Corning). 1X HBSS only sham controls were also utilized. Recipient mice were preconditioned by same-day sublethal irradiation (500 rads) and anesthetized via isoflurane (MWI Veterinary Supply Company) for injections. Peripheral blood was assessed as described above. Donor-derived chimerism was determined as the frequency of GFP-cells among a given cell population, with the baseline frequency background-subtracted. Similarly, cell number was determined via quantification of GFP- (donor-derived) cells as described above, background subtracted against the baseline number.

### HSC transplantations

HSCs from young and aged mice were freshly isolated via FACS from cKit-enriched BM as described above. Populations were sorted twice to ensure specificity. For each individual transplant, 2,500 HSCs were injected retro-orbital in 50μl 1X HBSS (Corning). Recipient mice were preconditioned by same-day sublethal irradiation (500 rads) and anesthetized via isoflurane (MWI Veterinary Supply Company) for injections. After 16-20 weeks of reconstitution, BM was harvested and assessed via flow cytometry as described above. Donor-derived chimerism was determined as the frequency of fluorescent cells among a given cell population. Cell numbers were similarly determined and as described above.

List of reagents and software can be found in Supplemental Table 3.

### Statistical analysis

Statistical tests are indicated in figure legends and performed using GraphPad Prism. For all tests, *p<0.05, **p<0.01, ***p<0.001, ****p<0.0001, and “ns” indicates non-statistical significance. Unless otherwise indicated, graphs of summary statistics indicate the mean ± standard error of the mean (SEM). The number of individual experiments and samples are outlined in the figure legends.

## Data availability

All sequencing data is available via the NCBI GEO Accession repository. Bulk RNAseq data was previously published^2^ (GSE226318). scRNAseq data was previously published^2^ (GSE255019). scCITEseq data is available via GSE280357. Note that the combined scRNA/CITEseq data can be accessed via the GSE280510 accession super series.

## Acknowledgments

We thank the University of California Santa Cruz flow cytometry (RRID SCR_021149), stem cell culture (SCR_021353), and vivarium facilities. This study was supported by the National Institutes of Health (NIH) National Institute on Aging (NIA) R01AG062879 to E.C.F., NIH National Institute of General Medical Sciences (NIGMS) K12GM139185 to B.A.M. and A.R.y.B., NIH NIGMS R25GM104552 to S.B.A., NIH Eunice Kennedy Shriver National Institute of Child Health and Human Development (NICHD) T32HD108079 to S.C., and California Institute for Regenerative Medicine (CIRM) EDUC4-12759 to M.G.E.R.

## Author contributions

B.A.M. and E.C.F. conceived of the study. B.A.M. wrote the manuscript with input from all authors. P.M. performed the bioinformatics analyses with input from L.M. and B.A.M. with oversight from V.D.J. B.A.M., S.S-B., A.R.y.B., E.B., A.D., S.B.A., S.C., M.R., and M.G.E.R. performed all experiments and data analysis. All authors reviewed and contributed to manuscript revisions.

## Declaration of interests

The authors declare no competing interests.

## List of abbreviations used

HSC: hematopoietic stem cell
MPP: multipotent progenitor
CLP: common lymphoid progenitor
CMP: common myeloid progenitor
GMP: granulocyte monocyte progenitor
MEP: megakaryocyte erythroid progenitor
EP: erythroid progenitor
MkP: megakaryocyte progenitor
cMkP: canonical megakaryocyte progenitor
ncMkP: non-canonical megakaryocyte progenitor
Tom: dtTomato
FACS: fluorescence activated cell sorting
MFI: median fluorescence intensity
BM: bone marrow
GM: granulocyte/monocyte
RBC: red blood cell

## Notes

### Competing Interest Statement

The authors have declared no competing interest.

