## Supplemental Data for "A rare HSC-derived megakaryocyte progenitor accumulates via enhanced survival and contributes to exacerbated thrombopoiesis upon aging"

Supplemental Figures 1-9
Supplemental Figure legends 1-9

See online version for Supplemental Tables 1-3.

#### Supplemental Figure 1

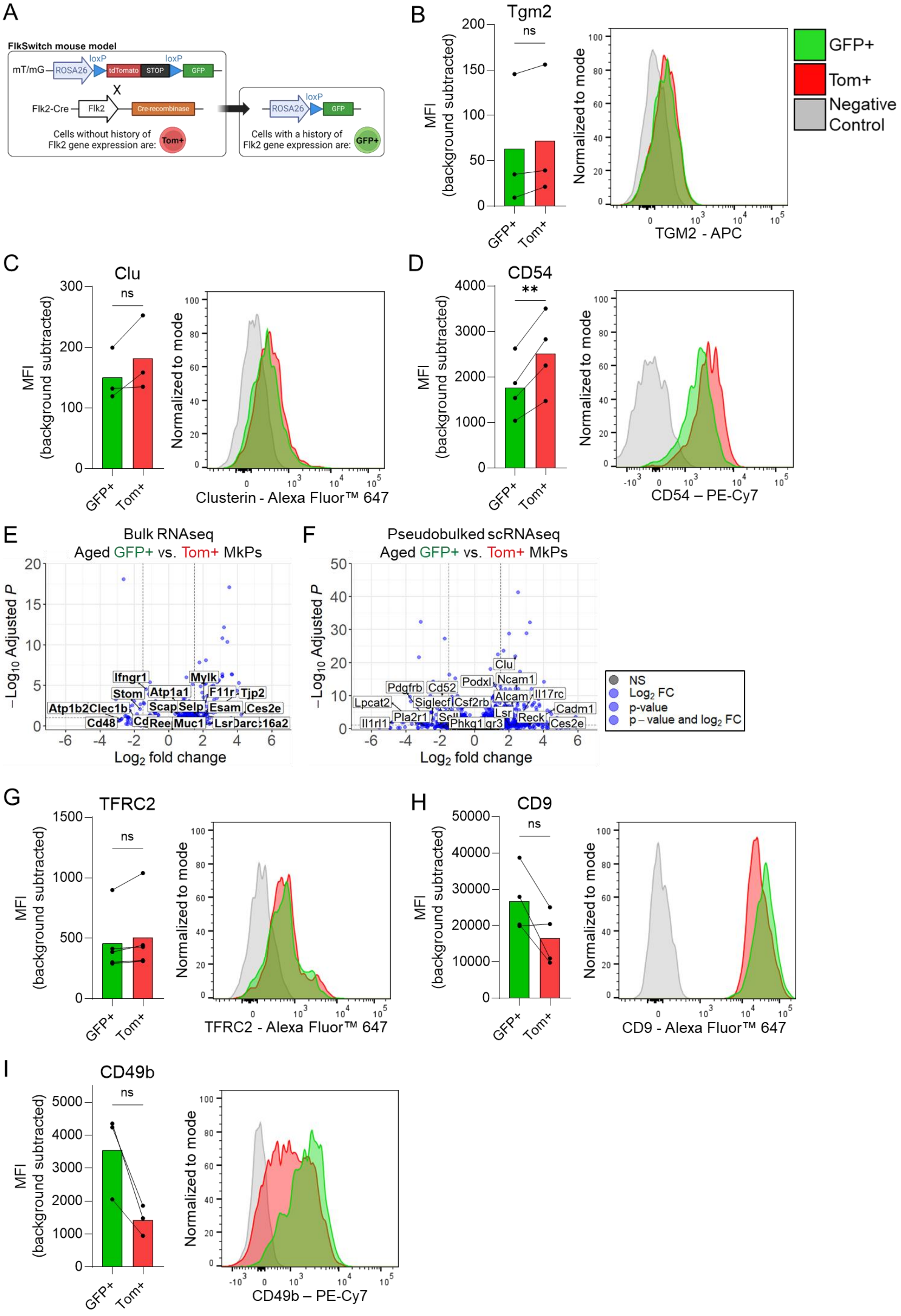

**Supplemental Figure 1. Transcriptomic and phenotypic heterogeneity of aged GFP+ and Tom+ MkPs.**

**A-C and F-G.** Flow cytometry analysis for the expression of the indicated marker on aged GFP+ and Tom+ MkPs from FlkSwitch mice. Each point represents an individual mouse with lines connecting values from the same mouse. Example histograms demonstrating staining pattern. n=3-5 across 6 independent experiments. **p<0.01 by paired t-test.

**D-E.** Volcano plots of **E.** bulk and **F.** scRNAseq data comparing aged FlkSwitch GFP+ and Tom+ MkPs from Poscablo et al., 2024[1]. Thresholds are adjusted p-value <0.1 or absolute $log_2$ fold change ≥1.5, indicated by blue points. Labeled genes indicate those predicted to encode cell surface proteins that have commercially available flow cytometry compatible antibodies. Only the top 10 genes in each direction by $log_2$ fold change displayed for graphical simplicity, excluding those indicated in Figure 1C-D. See Table 1 for complete gene lists. **D.** 8120 total genes and **E.** 11570 total genes plotted.

### Supplemental Figure 2

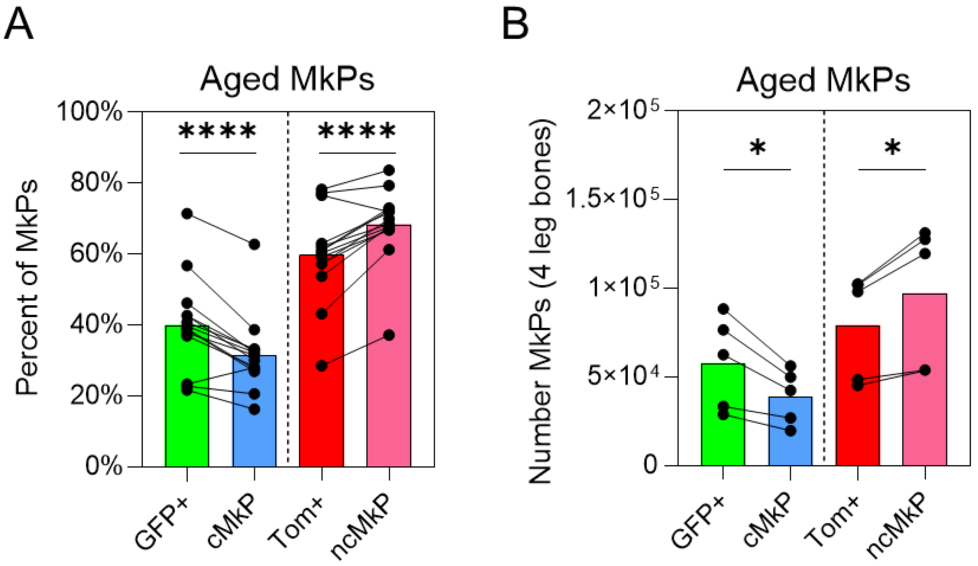

**Supplemental Figure 2. Inter-mouse assessment of aged cMkPs and ncMkPs.**

**A-B.** The **A.** frequency and **B.** number of aged MkP populations, comparing GFP+ to cMkP and Tom+ to ncMkP via paired analyses (same data as in Figure 2C-D). Each point indicates an individual animal with lines connecting the same mice. **A.** n=14 across 9 independent experiments and **B.** n=5 across 2 independent experiments. *$p<0.05$ and ****$p<0.0001$ by paired t-test.

### Supplemental Figure 3

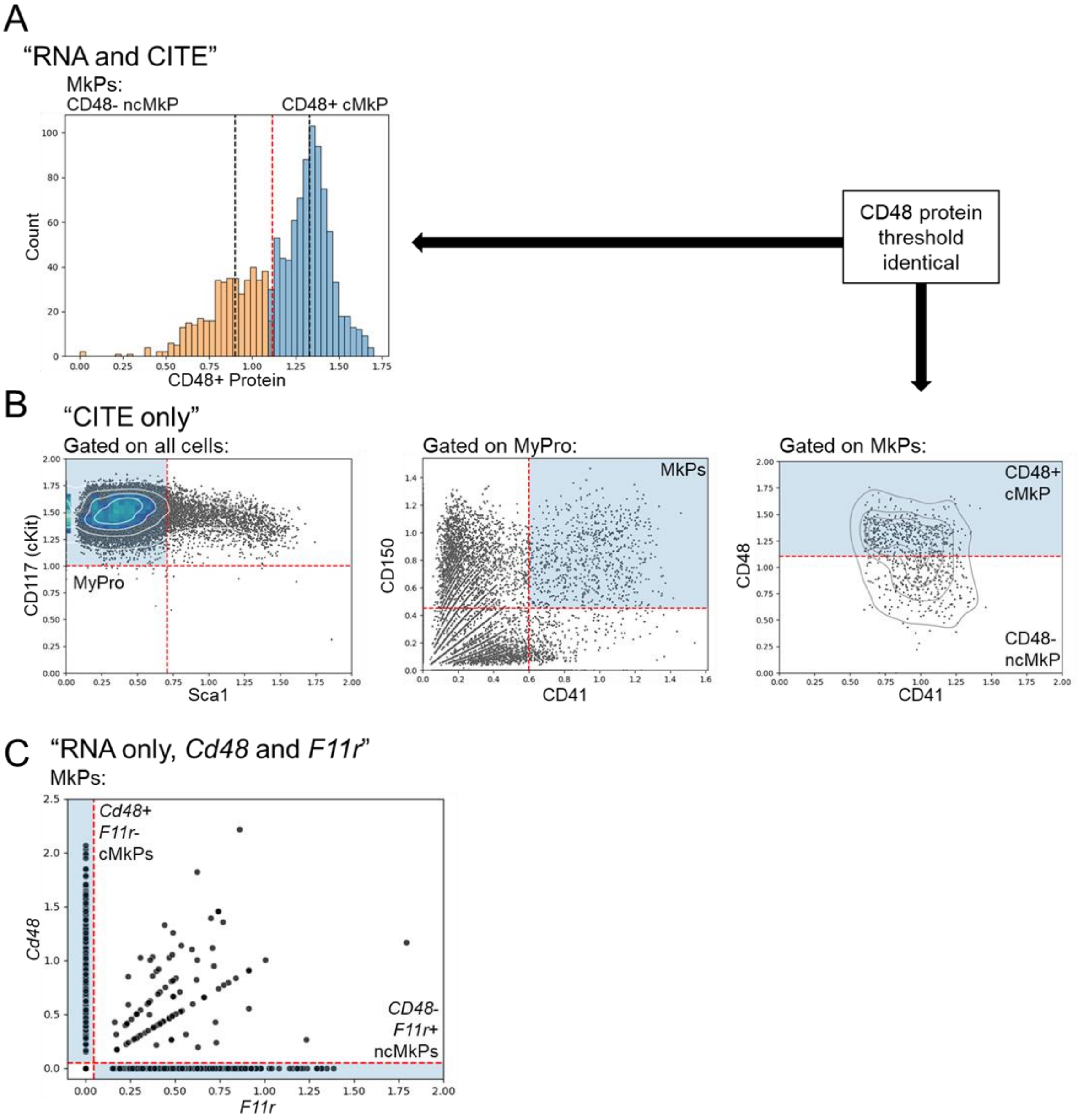

**Supplemental Figure 3. Additional scRNA/CITEseq analysis of aged FlkSwitch bone marrow MkPs.**

**A.** Histogram showing CD48 protein expression among all scRNAseq-annotated MkPs. Gaussian mixture model was used to classify MkPs as cMkPs and ncMkPs based on CD48 protein expression. Black dashed line is the median of the cMkP and ncMkP CD48 distributions, while the red dashed line is the mean of the two medians.

**B.** Protein gating strategy of scCITEseq data to annotate MyPro and MkP populations similar to flow cytometry. Red dashed lines and blue shaded boxes show thresholds used to annotate cells.

**C.** Annotation of cMkP and ncMkP via *F11r* and *Cd48* RNA expression among all scRNAseq annotated MkPs. *Cd48+F11r-* represent cMkPs whereas *Cd48-F11r+* indicate ncMkPs. Red dashed lines and blue shaded boxes indicate thresholds used to annotate cells.

**D.** Diffusion pseudotime analysis of aged MkPs from the scRNA/CITEseq data demonstrate striking similarity. Each point represents a single cell. **p<0.01 and ***p<0.001 by one-way ANOVA adjusted for multiple comparisons via Tukey's test. GFP+ and cMkP groups compared against each other and Tom+ and ncMkPs similarly compared.

#### Supplemental Figure 4

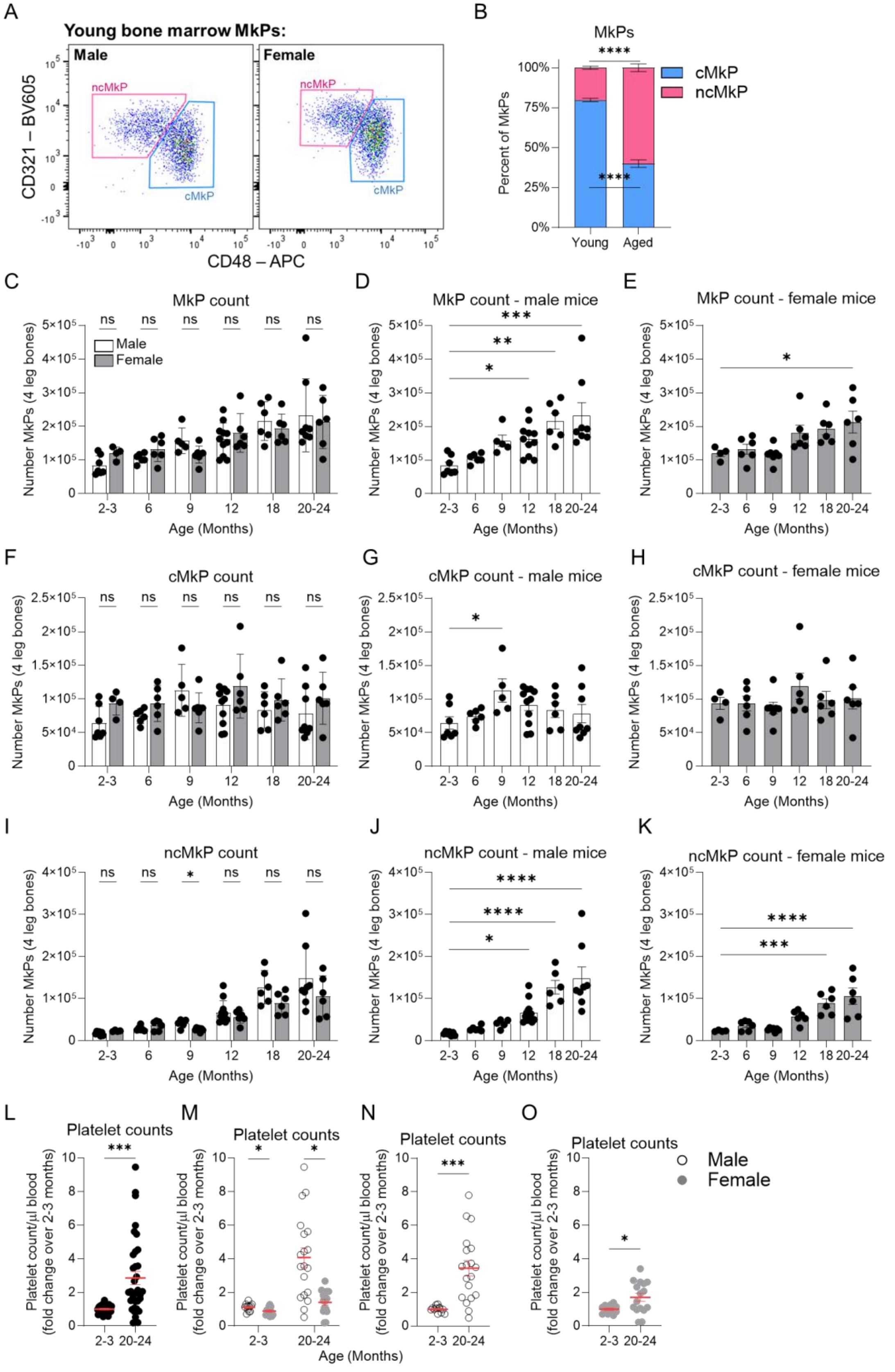

**Supplemental Figure 4. Assessment of megakaryopoiesis cell states throughout life.**

**A.** Example flow cytometry plot demonstrating detectable, yet rare, ncMkPs in young mice.

**B.** Comparison of the cMkP and ncMkP frequency between young and old WT mice. n=39 and old n=43 mice across more than 10 independent experiments. Unpaired t-test, ****$p<0.0001$.

**C-K.** Quantification flow cytometry analysis of BM MkPs from male and female mice. Each point represents an individual mouse. n=11, 12, 12, 17, 12, and 14 for ages 2-3, 6, 9, 12, 18, and 20-24 months, respectively across 10 independent experiments.

**C, F, I.** Comparison of **C.** total MkPs, **F.** cMkPs, and **I.** ncMkPs between male and female mice across six age groups. *$p<0.05$ by multiple unpaired t-tests with Holm-Šídák multiple comparisons adjustment.

**D-E, G-H, J-K.** Comparison of male and female **D-E.** total MkPs, **G-H.** cMkPs, and **J-K.** ncMkPs numbers over time. *$p<0.05$, **$p<0.01$, ***$p<0.001$, and ****$p<0.0001$ by one-way ANOVA, comparing all groups to the 2-3 month age group by Dunnett's multiple comparisons test.

**L-O.** Total platelet counts/μl blood as determined by quantitative flow cytometry, displayed as fold change over 2-3 month old mice. Each point represents an individual mouse and error bars are mean ± SEM. n=24 young and n=35 old across 5 independent experiments.

**L, N-O.** *$p<0.05$ and ***$p<0.001$ by unpaired t-test.

**M.** *$p<0.05$ by multiple unpaired t-tests with Holm-Šídák multiple comparisons adjustment.

#### Supplemental Figure 5

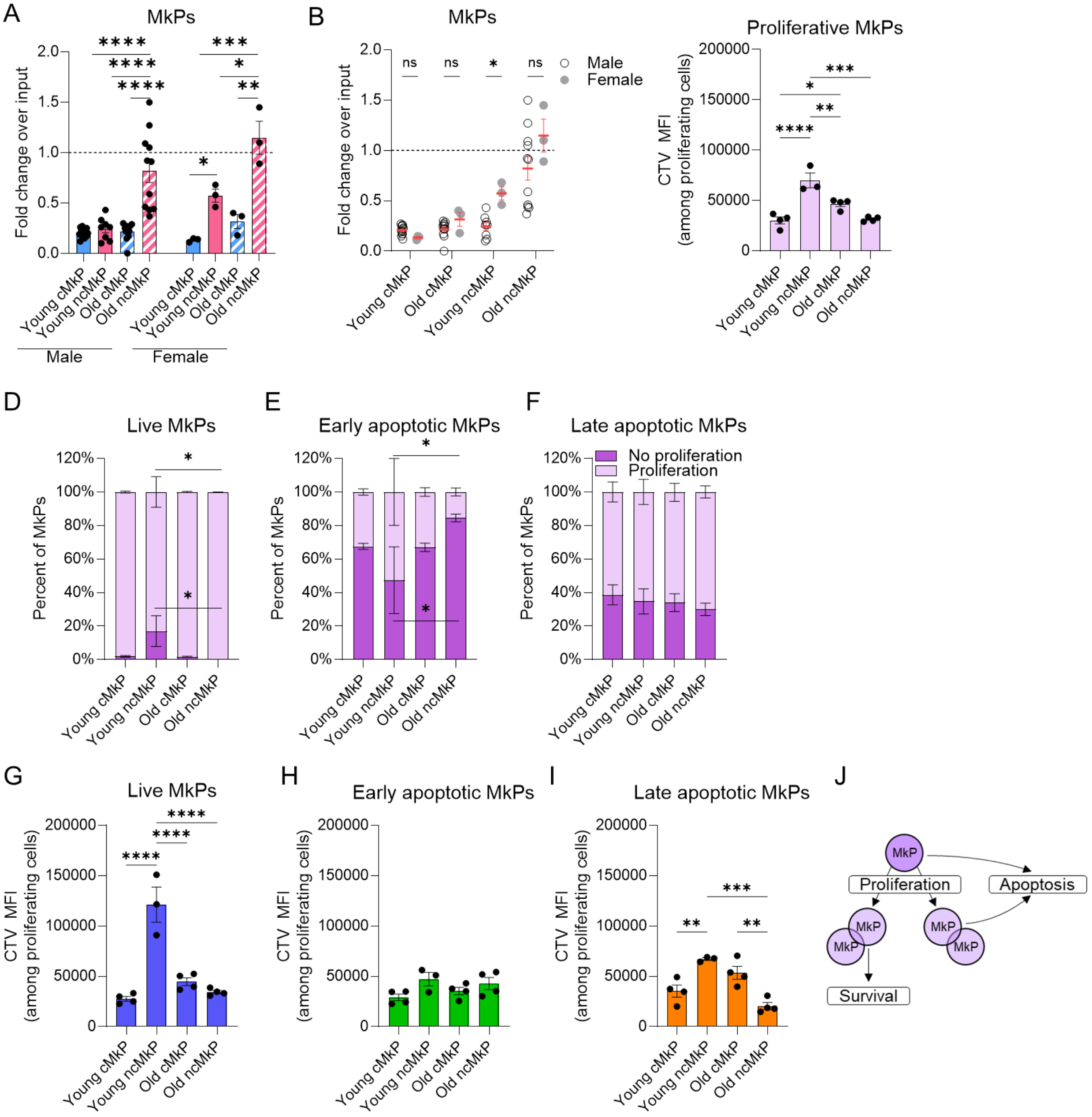

**Supplemental Figure 5. ncMkPs have a survival advantage in vitro.**

**A-B.** Comparison within **A.** or between **B.** MkPs from male and female mice. n and experimental design as in Figure 4A.

**C.** MFI of CTV among proliferating MkPs.

**D-F.** Proportion of in vitro cultured MkPs that proliferated over the three day culture, stratified by viability cell state.

**G-I.** MFI of CTV among proliferating MkPs of the indicated viability states.

**C-I.** n=3-4 across 4 independent experiments. Up to three technical replicates per mouse were averaged together as cell number allowed.

**J.** Model of in vitro MkP proliferative and survival performance.

*p<0.05, **p<0.01, ***p<0.001, and ****p<0.0001 by one-way ANOVA adjusted for multiple comparisons via Tukey's test or multiple unpaired t-tests (similarly adjusted for multiple comparisons).

**Supplemental Figure 6**

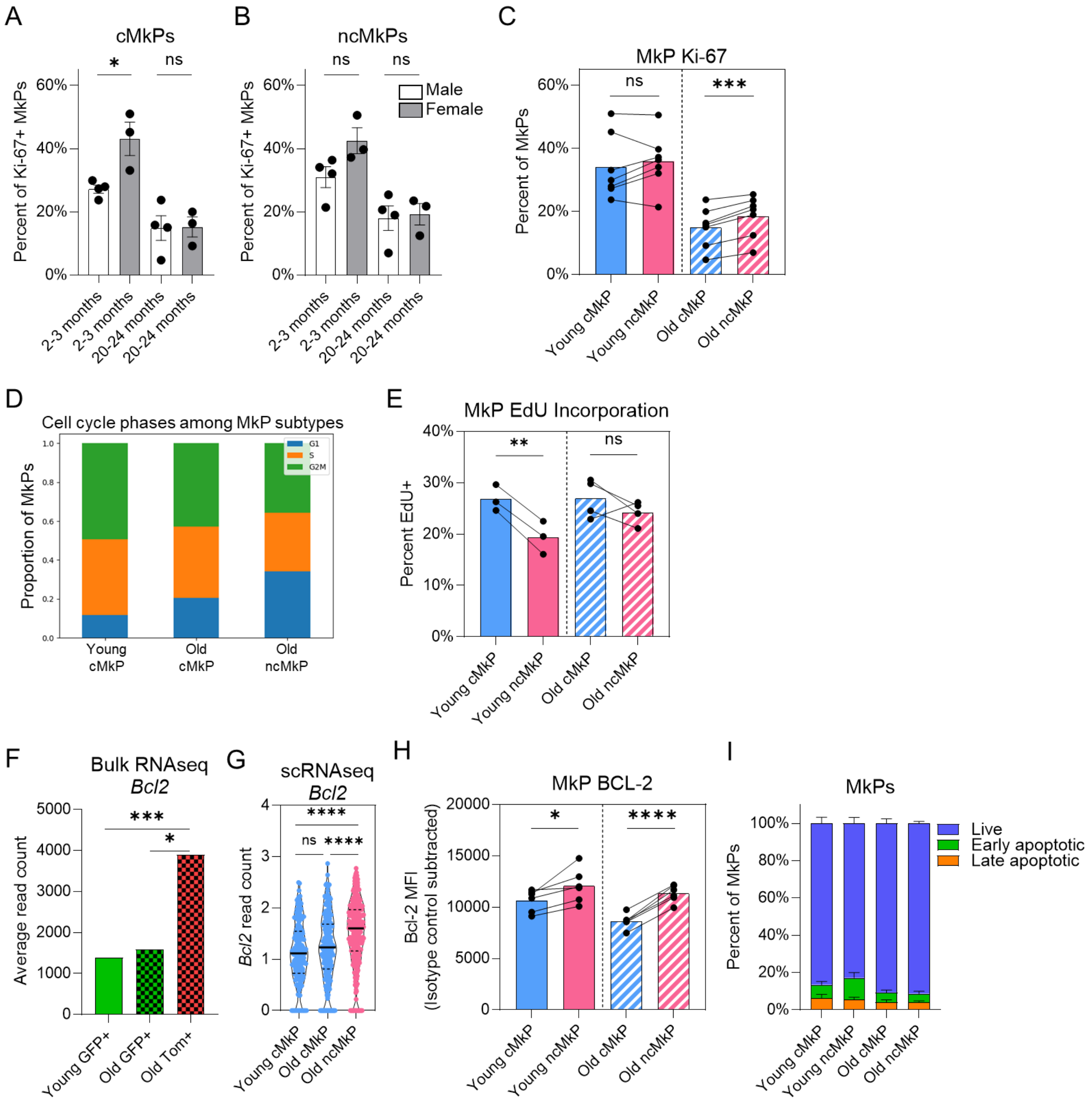

**Supplemental Figure 6. In situ features of MkP subtypes.**

**A-C.** Frequency of Ki-67+ cells among young and old MkP subtypes from freshly isolated BM. **A-B.** Comparison between male and female mice. *p<0.05 by unpaired t-test. **C.** Comparison within mice. Each point represents a single mouse with lines connecting cells from the same mouse. ***p<0.001 by paired t-test. n and experimental design as in Figure 5A.

**D.** Proportion of MkP cell cycle phases inferred from scRNAseq signatures. Young ncMkPs omitted due to low cell numbers.

**E.** Comparison of 24 hour EdU incorporation within individual mice. Each point represents a single mouse with lines connecting cells from the same mouse. **p<0.01 by paired t-test. n and experimental design as in Figure 5B.

**F-G.** Read count data of *Bcl2* from **F.** bulk and **G.** scRNAseq. Statistical significance determined via DESeq2 comparisons[1,2] for **F.** and *p<0.05, ***p<0.001, and ****p<0.0001 by one-way ANOVA adjusted for multiple comparisons via Tukey's test for **G.**

**H.** Paired analysis within individual mice for relative abundance of BCL-2 protein. *p<0.05 and ***p<0.0001 by parried t-test. n and experimental design as in Figure 5C.

**I.** Proportion of viability cell states among freshly isolated MkPs. n=6-8 across 4 independent experiments. Statistical non-significance determined by one-way ANOVA adjusted for multiple comparisons via Tukey's test, each cell state tested separately.

#### Supplemental Figure 7

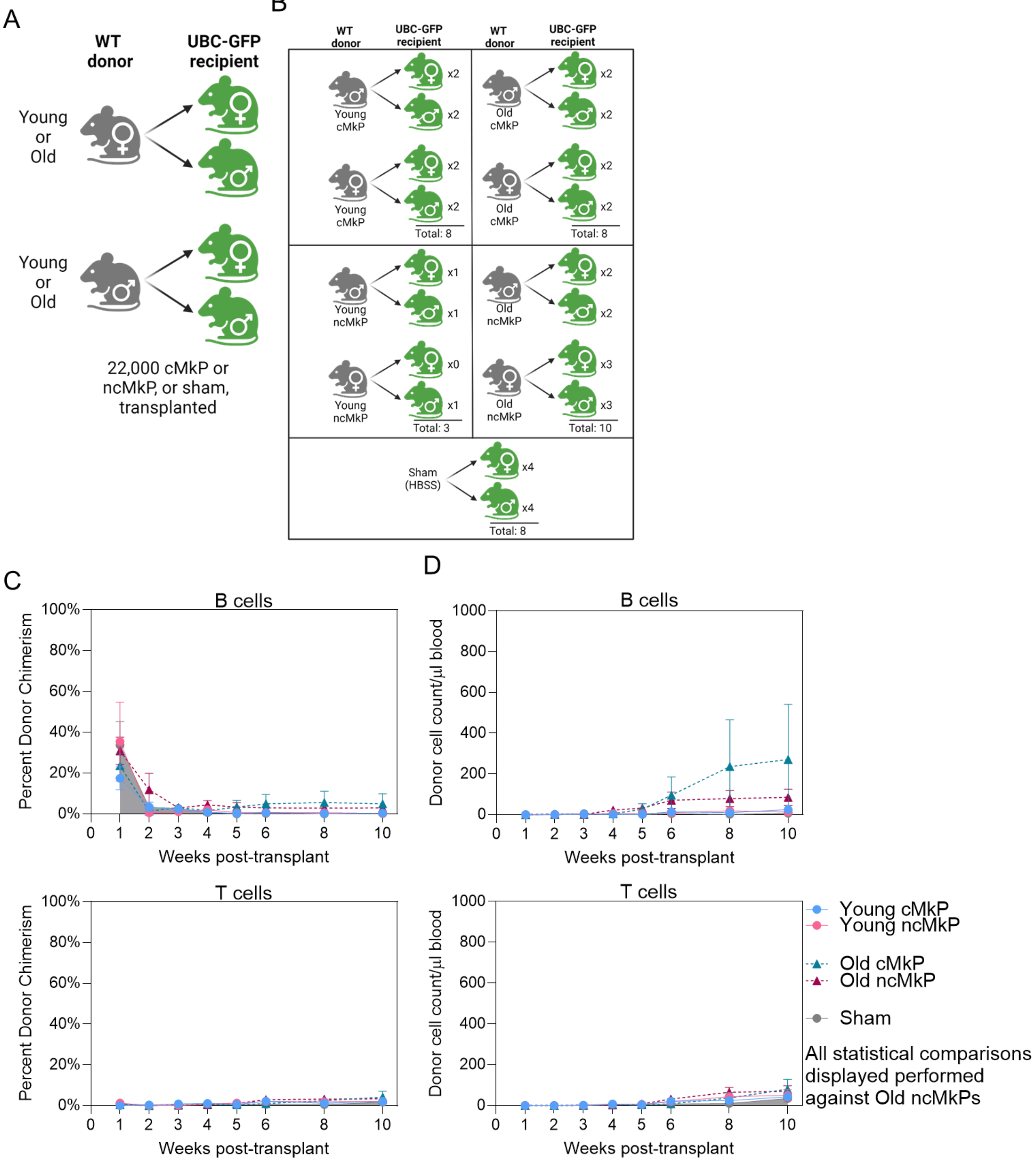

**Supplemental Figure 7. MkP transplantations and SDF1 migratory capacity.**

**A-B.** MkP transplantation schematic and design.

**C-D.** The percent donor chimerism **C.** and donor-derived cell number **D**. of the indicated populations. n, experimental design, and statistics as in Figure 5E-F.

#### Supplemental Figure 8

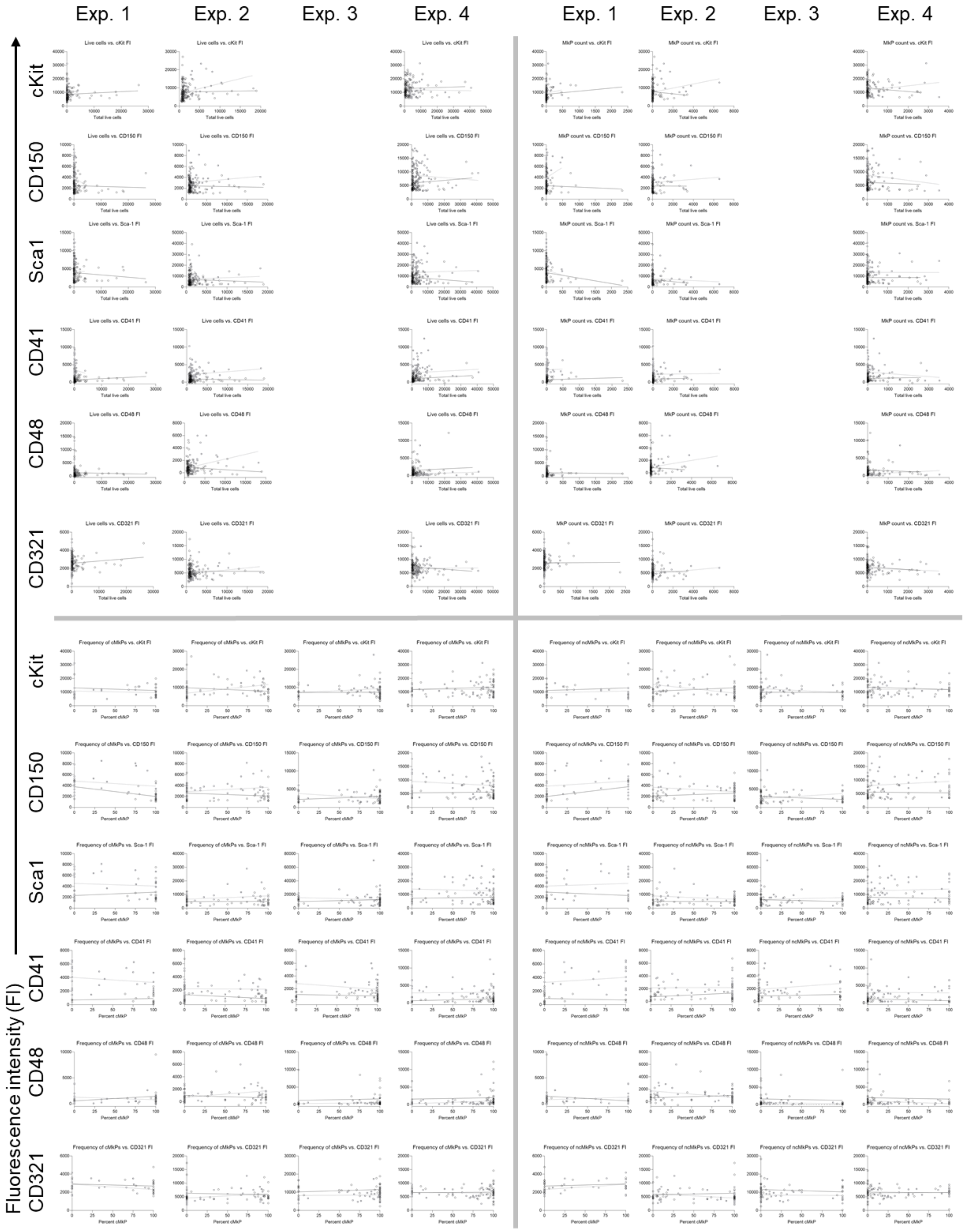

**Supplemental Figure 8. HSC phenotype does not correlate with MkP output in vitro.**

Flow cytometry single index sorted HSCs were correlated against their live cell and MkP output, comparing fluorescence intensity of each indicated marker against cell number or percent cMkP output across four experiments. Young are black open circles and trend line, whereas filled grey circles and trend line represent aged HSCs. See Supplemental Table 2 for statistical report (Spearman two-tailed correlation). n and experimental design are as in Figure 7.

### Supplemental Figure 9

A

| Cell population | Abbreviation | Phenotypic definition |
|---|---|---|
| cKit+Lineage$^{low}$Sca1+ | KLS | Lineage$^{low}$, cKit+Sca1+ |
| Hematopoietic stem cell | HSC | Lineage$^{low}$, cKit+Sca1+CD150$^{high}$Flk2-<br>or<br>Lineage$^{low}$, cKit+Sca1+CD150$^{high}$CD48- |
| Myeloid progenitor | MyPro | Lineage$^{low}$, cKit+Sca1- |
| Megakaryocyte progenitor | MkP | Lineage$^{low}$, cKit+Sca1-CD150+CD41+ |
| Platelet | N/A | CD11b-Gr1-B220-CD3-Ter119-CD41+ |
| Red blood cell | RBC | CD11b-Gr1-B220-CD3-Ter119+CD41- |
| Granulocyte/monocyte | GM | Ter119-B220-CD3-CD11b+Gr1+ |
| B cell | N/A | Ter119-CD11b-Gr1-CD3-B220+ |
| T cell | N/A | Ter119-CD11b-Gr1-B220-CD3+ |
| All populations pregated on single, live cells unless otherwise indicated. Specific subpopulations as indicated in the text and figures. | | |

B

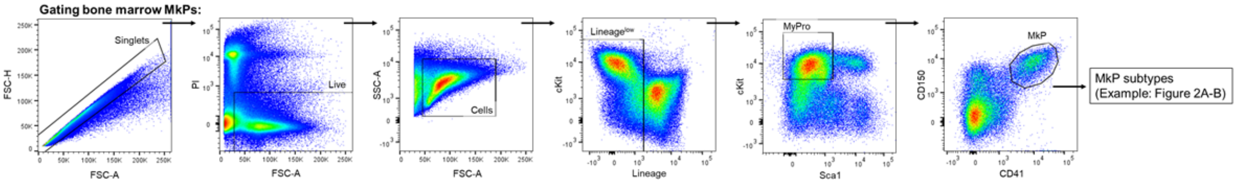


C

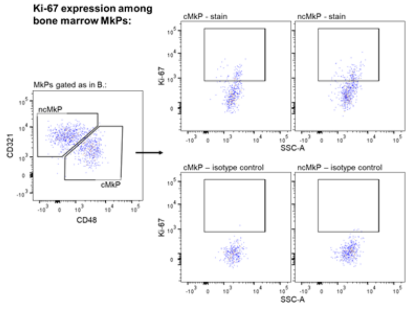


D

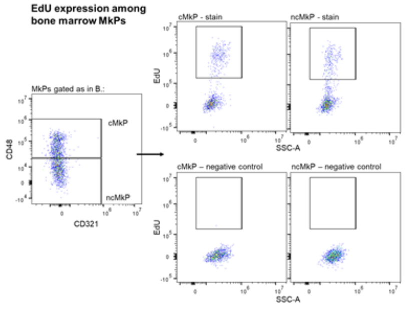


E

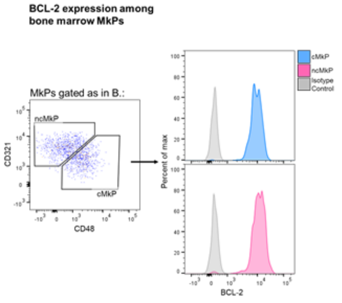

### Supplemental Figure 9 continued

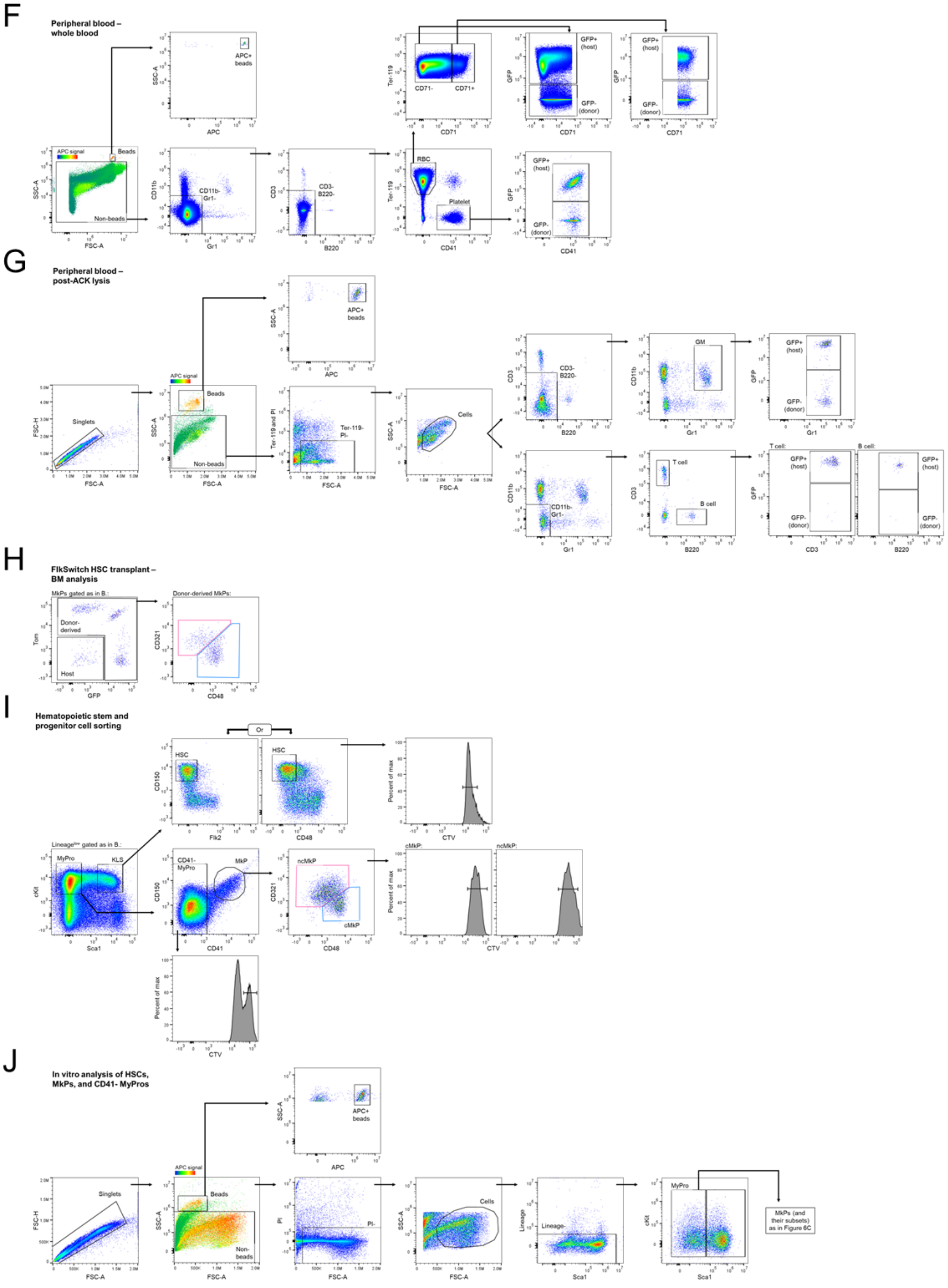

**Supplemental Figure 9. Phenotypic cell definitions and representative flow cytometry gating strategies.**

**A.** Table of all phenotypic cell definitions used.

**B.** Representative sequential flow cytometry gating strategy of BM MkPs.

**C.** Example intranuclear staining of BM MkPs for Ki-67 expression. The frequency of Ki-67+ cells (top row) for any given population was background subtracted using the isotype control (bottom row).

**D.** Assessment of EdU incorporation by BM MkPs. Not that the experimental procedure ablates CD321 expression, thus cMkPs and ncMkP defined by CD48 expression alone. The frequency of EdU+ cells (top row) for any given population was background subtracted using the negative control (bottom row).

**E.** Representative intranuclear assessment of BCL-2 among BM MkPs. The MFI of BCL-2 was background subtracted against the isotype control following standardized flow cytometry practices[3].

**F-G.** Example sequential staining of peripheral blood, either as **F.** whole blood or **G.** following ACK lysis. Detection of APC+ Calibrite beads allows quantitation of numbers of cells/μl blood (see Methods). Determining donor chimerism of each population in MkP transplantations as shown in plots assessing GFP positivity.

**H.** Assessment of frequency and number of donor-derived cMkPs and ncMkPs following HSC transplantations.

**I.** Example FACS plots for HSCs, cMkPs, and ncMkPs. When in vitro CTV experiments were performed (see Methods), only uniform CTV+ cells were sorted as indicated.

**J.** Example in vitro analysis of HSCs, CD41- MyPros, or MkP subpopulations. Sequential staining and bead-based determination of cell number identical for all approaches.
